# Cell motility greatly empowers bacterial contact weapons

**DOI:** 10.1101/2023.10.10.561656

**Authors:** Sean C. Booth, Oliver J. Meacock, Kevin R. Foster

## Abstract

Many bacteria kill competitors using short-range weapons, such as the Type VI Secretion System (T6SS) and Contact Dependent Inhibition (CDI). While these can deliver powerful toxins, they rely on direct contact between attacker and target cells. We hypothesised that movement enables attackers to contact more targets and thus greatly empower their weapons. To explore this, we developed individual-based and continuum models to show that motility greatly improves contact-dependent toxin delivery through two underlying processes. First, genotypic mixing increases the inter-strain contact probability of attacker and sensitive cells. Second, target switching ensures attackers constantly attack new cells, instead of repeatedly hitting the same cell. We test our predictions with the pathogen Pseudomonas aeruginosa, using genetically engineered strains to study the interaction between CDI and twitching motility. As predicted, we find that motility massively improves the effectiveness of CDI, in some cases up to 10,000-fold. Moreover, we demonstrate that both mixing processes occur using timelapse single-cell microscopy and quantify their relative importance by combining experimental data with our models. Our work shows how bacteria combine cell movement with contact-based weapons to launch powerful attacks on their competitors.

## Introduction

Many bacteria employ contact weapons against their competitors. These systems include the Type VI Secretion System (T6SS) that fires a toxin-laden needle into nearby cells, and Contact Dependent Inhibition (CDI) systems that deploy surface filaments to deliver toxins to adjacent cells after recognition of an outer membrane receptor (1). The strength of these systems is their ability to directly deliver a potent toxin into a competing cell, and there is growing evidence that contact weapons are often critical to whether a given strain succeeds in its preferred niche (2–4). However, they also suffer from a major limitation. Their short range means that an attacker can only target cells in its immediate vicinity. If all adjacent cells are clonemates or competing cells that have already been killed, the weapons will have no effect (5–7).

Bacteria actively move through their environments, which can be critical for their ability to compete for territory and colonise hosts (8, 9). Using a variety of different molecular appendages, they move by sliding, gliding, twitching and swimming (10). However, it is surface-associated motility mechanisms such as gliding motility or Type IV pilus-based twitching motility that are most commonly used when cells are at the high cell densities where contact-based weapons are effective (11, 12). Consistent with this, these conditions are also closely associated with the expression of T6SS (13) and CDI genes (14). Moreover, deleting genes for motility systems can reduce the impact of contact weapons (15–17) suggesting that motility systems impact close-range combat.

Based on these observations, we hypothesized that motility may serve to greatly improve the functioning of contact-based weapons by allowing attacker cells to reach more target cells. In order to explore this hypothesis, we developed an individual-based model that combines realistic cell movement in the crowded conditions of surface-attached communities (18) with contact warfare, and used a complementary continuum model to discern the underlying physical mechanisms. These models predict that cell movement can greatly improve contact weapons. Moreover, they identify two contributory processes: genotypic mixing and target switching. We test these predictions experimentally using genetically engineered strains of *Pseudomonas aeruginosa*, and find that, as predicted, twitching motility greatly improves the functioning of CDI during bacterial warfare. By studying competitions across a range of initial cell densities, we are also able to resolve the relative contributions of genotypic mixing and target switching during competitions. This analysis reveals that both processes help to explain the benefits of motility, but their relative importance can vary greatly as a function of initial conditions. Overall, our work shows how bacteria can combine motility with contact weapons as a highly effective strategy to overcome competitors.

## Results

### Individual-based modelling reveals two ways that motility empowers contact weapons

What is the role of motility in contact-dependent warfare? To explore this question, we developed a model that combines surface motility and contact weaponry. The strength of a modelling approach is that it is general and allows us to explore our question for any species of bacteria that displays surface motility and contact weaponry. Moreover, with models one can systematically and independently vary key properties, such as the level of motility and attack rate, on a scale that is not feasible experimentally. In this way, we can identify general principles that transcend a given experimental system. However, to demonstrate the validity of our model, we also use it to explain concrete experimental observations (below).

We adapted an Individual-Based Model (IBM) based on a pre-existing framework of cell motility (18, 19) in which each bacterial cell is simulated as a rigid rod that moves via a self-generated propulsion force *F* and pushes away neighbours (Methods). Cells can be one of two genotypes, either attacker or sensitive, and attacker cells deliver intoxification ‘hits’ to adjacent sensitive cells at a constant firing rate λ. All cells in a given simulation move with the same *F*, representing an intra-specific competition scenario in which motility is equivalent for attacker and sensitive strains. Our model is robust to changes in the microscopic intoxification mechanism, allowing us to encompass both T6SS-type (firing) and CDI-type (receptor-induced translocation) weapons with this framework (Methods, Fig. S1). Both attackers and sensitives are initially seeded into random locations at a seeding density *ρ*_0_, which are then expanded into clonal patches, replicating known growth patterns of mixed communities on surfaces (20, 21). The seeding density determines the level of initial spatial structure or ‘patchiness’, with lower seeding densities resulting in larger patches, which allows us to manipulate the initial degree of intermixing of the two strains. In our first simulations, we assess the efficacy of the weapon by tracking the rate at which sensitive cells receive their first hit, essentially modelling a toxin with single-hit killing kinetics (7). However, we will later consider toxins with multi-hit kinetics, whereby increasing numbers of hits have an increasingly strong impact on the growth of sensitive cells (22).

We used this model to identify the key processes that determine the effectiveness of contact weapons, i.e. the extent to which attacker cells are able to impact the survival of sensitive competitors. When the two genotypes are initialised in patches, a key limitation to non-motile attackers is that they are unable to come into contact with the majority of sensitive cells as their clonal patches remain separate. As a result, intoxification is limited to a one-cell thick region on the outside of the attacker patches (Fig. 1A,B, top row, Movie S1, top right). Introducing motility allows attackers to disperse this initial structure and encounter sensitive cells throughout their patches, substantially enhancing the efficiency of the attacking population (Fig. 1A,B, second row; Movie S1, top left).

**Figure 1.**
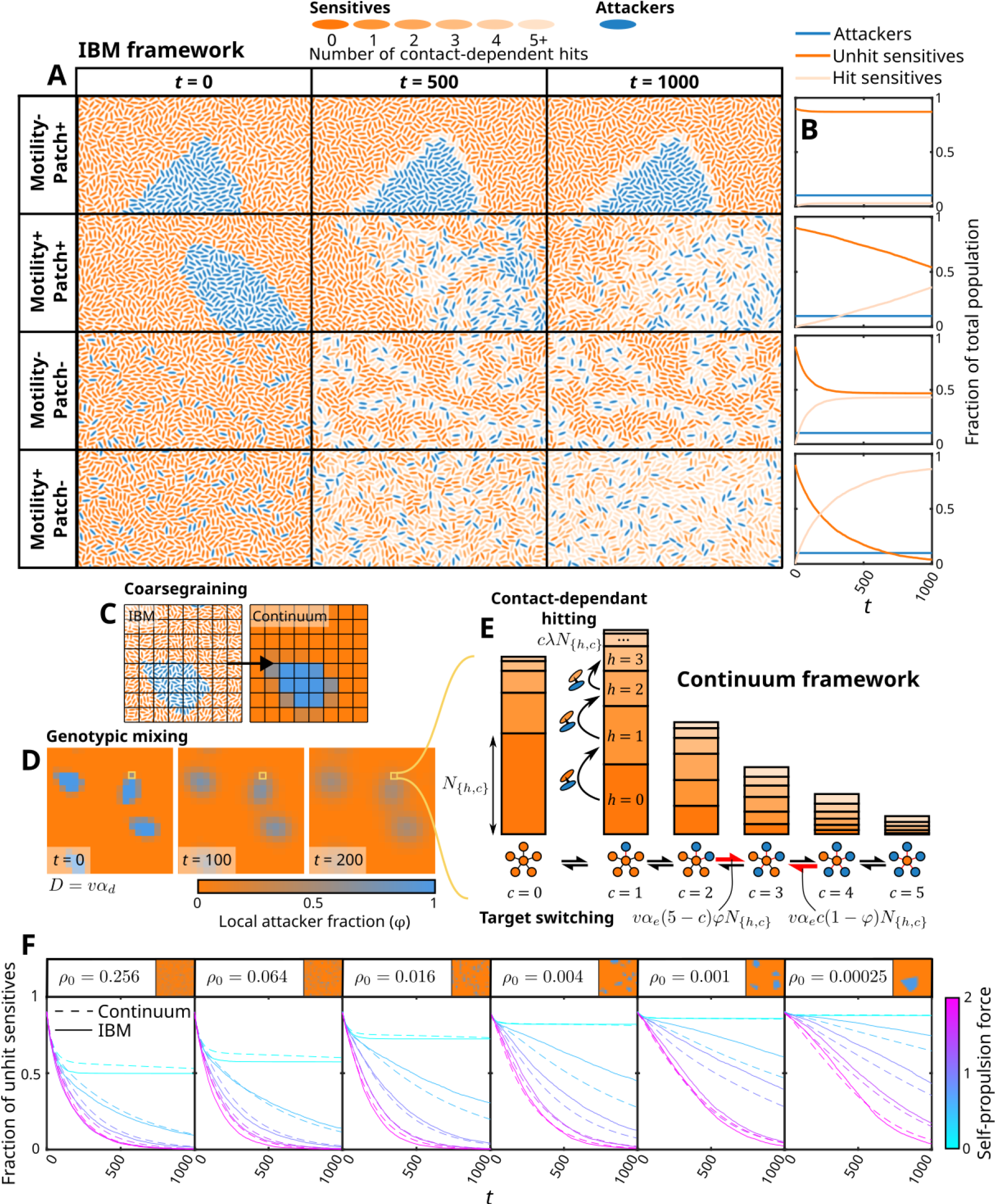
Individual-based and dynamical modelling approaches suggest three separate processes govern contact-dependant attack kinetics. (**A**) We used an Individual-Based Modelling (IBM) framework to allow simulation of contact-dependant hitting between separate attacker and sensitive populations under motile (motility +) and non-motile (motility –) conditions, as well as patchy (patch +) and non-patchy (patch –) starting population distributions. Increasing numbers of hits accumulated by sensitive cells are represented by their dark orange to light orange colouration.

Here, we consider a 1:9 attacker:sensitive ratio to illustrate the processes that impact intoxification dynamics. (**B**) Differences between the rates of hit accumulation under the different conditions suggest two separate processes determine competitive outcomes: large-scale diffusive mixing of inhomogeneous populations (‘genotypic mixing’) and small scale mixing of cell contacts (‘target switching’). (**C-E**) To test our understanding of this system, we constructed a continuum spatial model in which only three processes were explicitly invoked: genotypic mixing, (governed by the diffusion constant *D* given by the average cell velocity 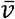 and a proportionality constant α_d_), target switching, (governed by the average cell velocity 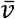, local attacker fraction *ω* and a proportionality constant *α*_*e*_) and contact-dependant hitting (governed by the firing rate λ). The top level of this model is built on a coarse-grained lattice representing the distribution of genotypes (**C**) on which diffusive genotypic mixing acts to gradually disperse any initial population structure (**D**). At a given lattice site (yellow square), we denote the fraction of the sensitive population contacting *c* attackers and hit h times as *N*_{*h,c*}_. These are represented in (**E**) using stacked bar charts; populations transition between different numbers of contacted attackers (x-axis) via target switching, and migrate to higher number of hits (y-axis, lighter shades of orange denote larger numbers of hits) via contact-dependant hitting. (**F**) By extracting *α*_*e*_ and *α*_*d*_ from the statistical properties of our IBM under different self-propulsion forces (Fig. S2, Supplementary Note 2), we arrive at a prediction of the behaviour of the IBM dynamics that contains no free parameters. The intoxification dynamics of the continuum model match that of the IBM across self-propulsion forces and seeding densities *P*_0_.

This effect, which we call ‘genotypic mixing’ here, was expected to have a strong effect on the intoxification dynamics as the degree of genetic intermixing determines the inter-population interaction frequency, making it a key factor in social phenotypes (17, 20, 23). However, even when we begin the model with the two genotypes fully intermixed, we find that intoxification is substantially more efficient when motility is active compared to when it is not (Fig. 1A,B, bottom two rows; Movie S1, bottom row). Inspection of the simulations reveal that this benefit arises by disrupting a process recently called the ‘corpse barrier’ effect (7), whereby stationary attackers are only able to hit a small group of sensitive neighbours which then act as shields for unhit sensitive cells behind them. As a result, attacker efficiency is greater when combined with motility because attacker cells can continuously switch out intoxified neighbours for fresh sensitive targets. We thus identify a second process independent of genotypic mixing – ‘target switching’ – by which motility empowers contact weapons.

In summary, our first model reveals two separate, but related, processes. First, *genotypic mixing* disrupts large-scale population structure and allows attackers to invade clonal patches of sensitives. Second, *target switching* disrupts corpse barriers and allows attackers to access fresh targets behind intoxified cells.

### Disentangling the two processes: Continuum modelling

The IBM identified two separate mixing mechanisms in our system. However, this model is limited in its ability to assess their relative importance for intoxication dynamics, or indeed whether additional processes are at play as the mixing processes are emergent properties of the IBM. To address this, we adopted an approach inspired by methods in theoretical physics. In statistical mechanics, the unpredictable behaviours of large numbers of particles can be shown to average out over large time and length scales, giving rise to predictable emergent phenomena such as diffusion (24). Here, we develop a coarse-grained description of our IBM in which we assume a similar large-scale averaging out of unpredictable behaviours occurs, allowing us to use deterministic descriptions of these emergent behaviours. By analogy, we assume that the unpredictable motion of the ‘particles’ in this model – the individual cells of the IBM – results in predictable macroscopic outcomes such as the winner of the competition between the two bacterial strains.

This approach provides two major advantages compared to using the IBM alone: firstly, by appropriately matching the coarse-grained description of the system to the microscopic dynamics of the individual IBM cells, we can test whether genotypic mixing, target switching, and contact-dependent firing are sufficient to explain the observed patterns in the IBM data. This allows us to confirm that we have not neglected a major, unknown contributor to the intoxification dynamics. Secondly, in contrast to the IBM, we can manipulate genotypic mixing and target switching independently in the continuum framework, allowing us to assess their relative importance in a given competition scenario.

We model the location of our sensitive and attacker populations on a lattice grid using the 2D phase-field *ω*(***r***), specifying that at locations ***r*** fully occupied by the sensitive population *φ*(***r***) = 0 and at locations fully occupied by the attacker population *φ*(***r***)= 1 (Fig. 1C). Genotypic mixing is then simulated by applying the diffusion equation to this phase field,

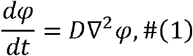

where *D* is the diffusion constant of the system, driven by cell motility, and *∇*^2^ is the 2D Laplace operator, which determines how the spatial structure of the system relates to diffusive concentration changes. Over time, the initial patchiness is smoothed out (Fig. 1D), reflecting the large-scale genotypic mixing of the two genotypes. The initial spatial structure of the attacker and sensitive populations is determined using a similar algorithm to that used in the IBM, allowing the two models to be directly compared when initialised with identical seed cell distributions.

Each lattice site ***r*** is treated as a well-mixed compartment, containing sub-populations *N*_{*h,c*}_ of sensitive cells that are labelled by the number of intoxification hit events *h* they have received and the number of attacking cells they are in contact with *c* (Fig. 1E) (e.g. *N*_{0,0}_ zero hits, in contact with zero attackers; *N*_{1,2}_ one hit, in contact with two attackers). The sum of all the sub-populations at a lattice site is equal to the total number of sensitive cells at that site, *i*.*e*. *Σ*_*h*_ *Σ*_*c*_*N*_{*h,c*}_(***r***) = 1 − *φ*(***r***). Sensitive cells thus exist in ‘states’ describing the number of hits accumulated and number of adjacent attackers, and cells can transition between states by increasing or decreasing the number of attackers they are in contact with and by increasing the number of hits they have accumulated. We model these state transitions as Markovian (i.e. transition rates depend solely upon the current system configuration), allowing us to simulate the population dynamics at lattice site ***r*** using the master equation

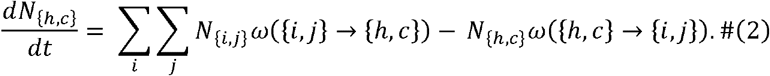

Here, the summation occurs over all possible states in the system {*i,j*}and the notation *ω*({1}→{2}) denotes the rate at which cells in state {1} transition to state {2}. The first term in this summation thus represents the rate at which cells in other states ‘jump into’ the focal state {*h,c*}, while the second represents the rate at which those cells in state {*h,c*} jump into other states. For example, the population in state [*h,c*} jumps out to {*h*+1,*c*} (i.e. accumulates a single hit) at a rate *ω*({*h,c*}*→* {*h* +1,*c*}) = cλ, which appears in the second term and reflects the dependence of the hit accumulation rate on both the single attacker cell firing rate λand the number of attacker cell contacts. On the other hand, the impossibility of losing hits is represented by a rate of zero for the inverse process, which appears in the first term as *ω*({*h*+1,*c*}→{*h,c*}) = 0. An explicit form of this master equation, in which the target switching rate *r*_e_ and local attacker fraction *φ*(***r***) appear in the terms determining the rate of change of the number of attacker contacts, is derived in Supplementary Note 1.

From Eqs. 1 and 2, we can deduce that the dynamics of the continuum model are determined by the rates of genotypic mixing (determined by *D*), contact-weapon firing (*λ*) and target switching (*r*_e_). We measured the dependence of *r*_e_ and *D* on the motility speed of cells in the IBM using independent simulations run without intoxication (Supplementary Note 2, Fig. S2), allowing us to use the continuum framework to predict the individual-based killing dynamics with zero free parameters (Fig. 1F, Movie S2). The extremely close match between the continuum and individual-based dynamics across three orders of magnitude of seeding densities and a wide range of self-propulsion forces (*i*.*e*. cell speeds) demonstrates that these three processes are sufficient to explain almost all the intoxification dynamics in the IBM. In addition, the continuum model is tractable to analytical techniques, which allows us to derive several insights into general properties of our system (such as saturation of intoxification with increasing cell velocity, Fig. S3) in Supplementary Note 3.

With all processes accounted for, we can now use the continuum framework to analyse the relative importance of genotypic mixing and target switching for contact warfare (Fig. S4). These analyses show that target switching is most important when seeding density is high, while genotypic mixing plays a greater role with low seeding density. Intuitively, this results from the fact that at low seeding densities, clonal patches tend to be of a larger spatial scale and so genotypic mixing is crucial. In contrast, when the system begins in a near well-mixed state at high seeding densities, the main limitation to weapon use is the build-up of dead cells around an attacker (corpse barriers). Under these conditions, target switching becomes more important.

### Experiments link cell motility to the effectiveness of contact warfare

We next sought to test our prediction that cell motility empowers contact weapons. The bacterium *Pseudomonas aeruginosa* PAO1 engages in surface ‘twitching’ motility using Type IV pili (25) and carries multiple close-range weapons. These include two independent Contact Dependent Inhibition (CDI) systems (14, 26), and three Type VI Secretion Systems (T6SS), where one, HSI-1, is dedicated exclusively to delivering anti-bacterial effectors (27). From these, CDI was selected as a close-range weapon, and a sensitive strain was generated by deleting the entire CDI 1 operon (PA0040-PA0041a) (14). CDI was used as the regulation of the anti-bacterial T6SS HSI-1 of *P. aeruginosa* is complex and it often has low activity in competition assays (28). Attacker and sensitive strains with limited motility were constructed by deleting the gene for the retraction motor *pilU* in the wild-type and ΔCDI background (26). These mutants can retract their pili using PilT, but retraction force is much lower such that cells can achieve very little movement by twitching motility on 1.5% agar (29). Modifications to the twitching motility system can change the expression of other virulence-associated genes through changes in intracellular cAMP levels (30, 31), which raised the possibility that expression of CDI would be altered in motility mutants. We chose to focus on *pilU*, therefore, because it retains approximately WT levels of cAMP induction while nevertheless largely eliminating cell movement (32). We reasoned that this approach should limit the potential for pleiotropic effects via changes to the twitching motility system. In addition, we used RT-qPCR to confirm that CDI genes were not differentially expressed in the Δ*pilU* strain (Fig. S5).

Using matched pairs with identical motility, attacker and sensitive strains were co-cultured (Fig. 2). Strains were competed using the colony biofilm model where strains are mixed together then allowed to grow on an agar surface (33, 34). Images were taken after 48h of competition (Fig. 2A, S6A), showing that, as expected, non-motile colonies were much smaller than motile colonies. Fluorescence intensities also suggested that more sensitive cells survive when motility is inactive than when it is active. To quantify competition outcomes, portions of the colonies were scraped off and plated. The fold-change in the ratio of attackers:sensitives from the start to end (deemed competitive advantage) was then calculated (Fig. 2B, S6B-D). At both the center and the edge of colonies, motility often drove a significant increase in the advantage for the attacker with large effect sizes; we observed up to a 10,000-fold increase in competitive advantage associated with motility. Only when the attacker was started at a 1:10 disadvantage was there no significant difference between motile and non-motile competitions, and then only at the colony center. The improvement in contact killing with motile cells was maintained across a wide range of initial inoculum (seeding) densities (Fig. S6B-D).

**Figure 2.**
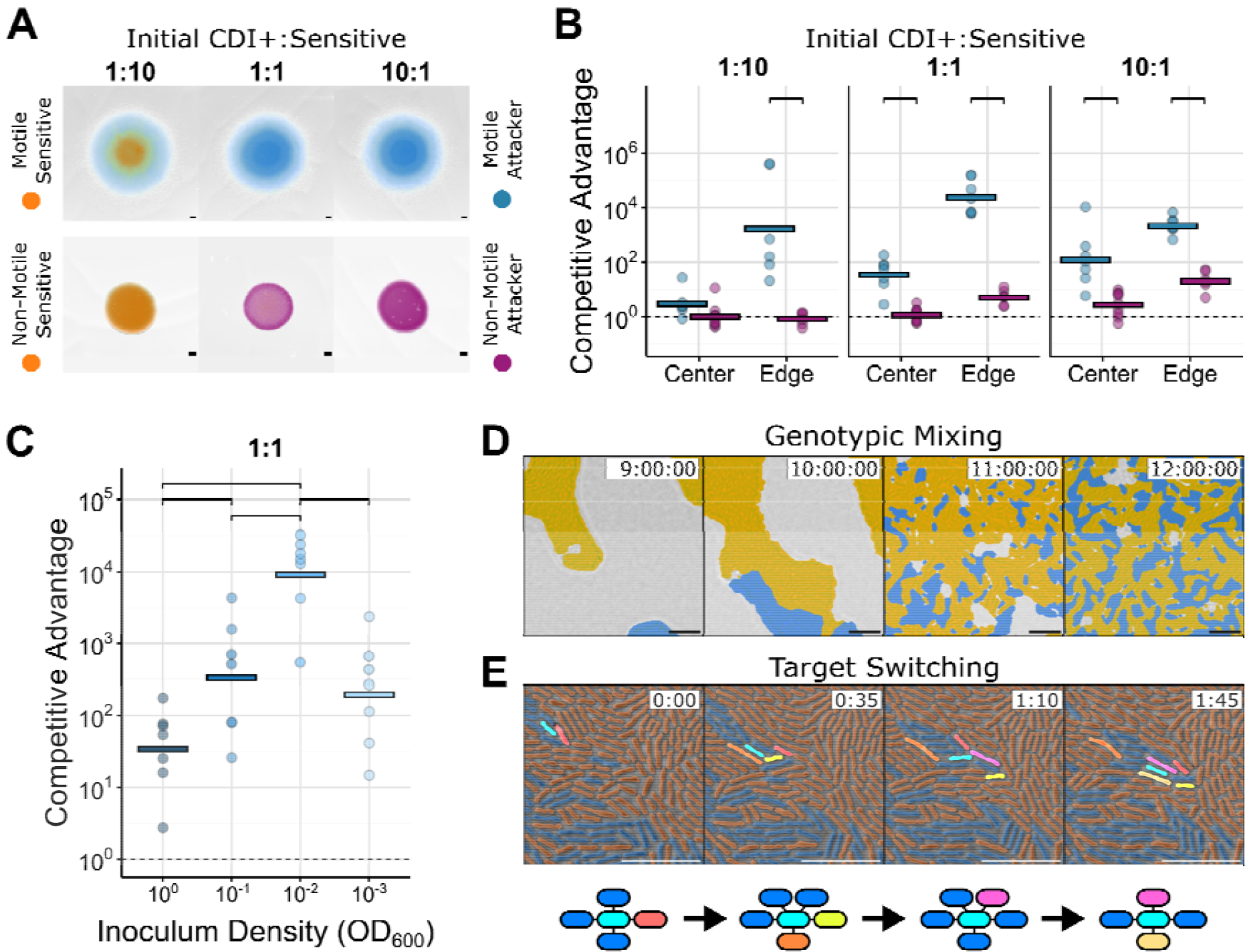
Experiments show that contact-dependent inhibition (CDI) is dependent on motility and is influenced by initial culture density. (**A**) Colony competitions were performed between wild-type or twitching motility deficient *P. aeruginosa* PAO1 and a corresponding engineered CDI-susceptible mutant. Colonies were inoculated with different initial ratios of attacker to susceptible cells. Representative microscopy images from motile (top) and non-motile (bottom) competitions after 48 h of growth show differences in the scale and structure of communities. (**B**) Quantification of the outcome of colony competitions from an inoculum density (OD600) of 1.0, picked at either the colony center or colony edge reveal a consistently large advantage for motile attackers. (**C**) However, quantification of motile colony competitions initiated from a range of inoculum densities (OD600 of 1.0, 0.1, 0.01 and 0.001) and equal ratio of attacker:susceptible cells demonstrate a non-monotonic relationship between inoculation density and the magnitude of this advantage. (D,E) Microscopy of colony dynamics reveals that both genotypic mixing (intermixing of the two strains from initial clonal patches, D) and contact switching (exchange of cells touched by an individual attacker, E) occur in this system. In A, strains are false-coloured either blue (motile CDI attacker), purple (non-motile attacker) or orange (motile and non-motile susceptible, top and bottom). Scale bars: A 500 μm; E, F 10 μm. In D, attacker cells are blue, sensitive are orange and unoccupied space is grey. In E, a focal attacker cell is highlighted in cyan. Sensitive cells newly contacted by this focal attacker in each frame are highlighted in unique colours. Networks below each timepoint indicate the changing collection of sensitive and attacker cells currently contacted by the focal attacker. Competitive advantage is calculated as the log fold-change in ratio of attacker:susceptible cells (as counted from sampling, plating and counting colony forming units) from the beginning to end of the experiment. Lines indicate the mean of replicates (n ≥ 6). Top brackets indicate a significant difference between densities (one-sided Welch’s t-test, p < 0.05, Benjamini-Hochberg MHT corrected 0.95). See supplementary Table 2 for exact group size (n) and p-values.

As discussed above, the models predict the existence of two different motility-dependent processes that determine the efficacy of contact-based weaponry. To assess whether these processes were occurring, we quantified cell motility over time in colonies inoculated with different initial densities by capturing short timelapses at single-cell resolution every 30 minutes (Movie S3). This confirmed the occurrence of both genotypic mixing (Fig. 2D) and target switching (Fig. 2E) in the experiments, reflecting the results of our models. In sum, as predicted by our theoretical approaches, our data show that motility greatly enhances the effectiveness of the CDI system.

### The importance of mixing and switching shift with initial conditions

While the general congruence between the model and data was encouraging, we sought a more robust test of the fit between our model and the data. One unexpected pattern in the data is that the relationship between inoculum density and competitive outcome in the motile strains is non-monotonic (*i*.*e*. shows an intermediate peak). Specifically, we observed in 1:1 competitions that the advantage provided by CDI initially increased with decreasing inoculum density. This pattern was unexpected as the initial degree of genotypic mixing scales linearly with inoculum density, meaning we observed an increased competitive advantage for CDI even as genotypic mixing was decreased. This advantage then dropped back down at the lowest tested density (Fig. 2C). This pattern is greatly weakened and no longer statistically significant in the experiments with the *Δ*pilU** motility mutant (Fig. S6D). Moreover, previous work with bacteria that lack surface motility also did not see this pattern, instead finding that higher inoculum density consistently enhances the efficiency of short-range weaponry (5, 17, 35, 36), likely due to the associated increase in initial intermixing of the two strains. These observations suggested that the non-monotonic relationship between inoculum density and CDI efficiency could be caused by changing effects of motility across densities.

To test this hypothesis, we parameterised our continuum model with cell movement data and asked if the model then recapitulates the non-monotonic relationship seen in the competition data. Specifically, we used the timelapse experiments described in the last section (Movie S4), to quantify cell motility in colonies inoculated with different initial densities. We focused on data from the colony centres, because this most closely resembles the conditions of the simulations where there is no outward expansion of the population as occurs at the colony edge. As cell velocity is a key input to our theoretical framework, we quantitatively measured the movement of large numbers of cells by using Particle Image Velocimetry (PIV) (37, 38), an image analysis method related to optical flow that tracks groups of pixels (Figs. 3B, S7B). Fluorescent images taken alongside each video (Figs. 3A, S7A) were then used to determine the percentage surface coverage by cells (Figs. 3C, S8A) and the extent of genotypic intermixing over time (Figs. 3E, S8C, Movie S5). For all densities, cell motility and genotypic intermixing increased until, through growth and division, the cells became confluent and filled the imaging window. Thereafter, the cells pushed into each other, velocity dropped, and 3D structures started to form. Lower cell densities resulted in higher cell velocities, with the highest velocities reached at the lowest two tested densities (Figs. 3D, S8B). The extent of intermixing only differed for the lowest initial density, as the three highest tested densities all ended up equally mixed (Fig. 3E).

**Figure 3.**
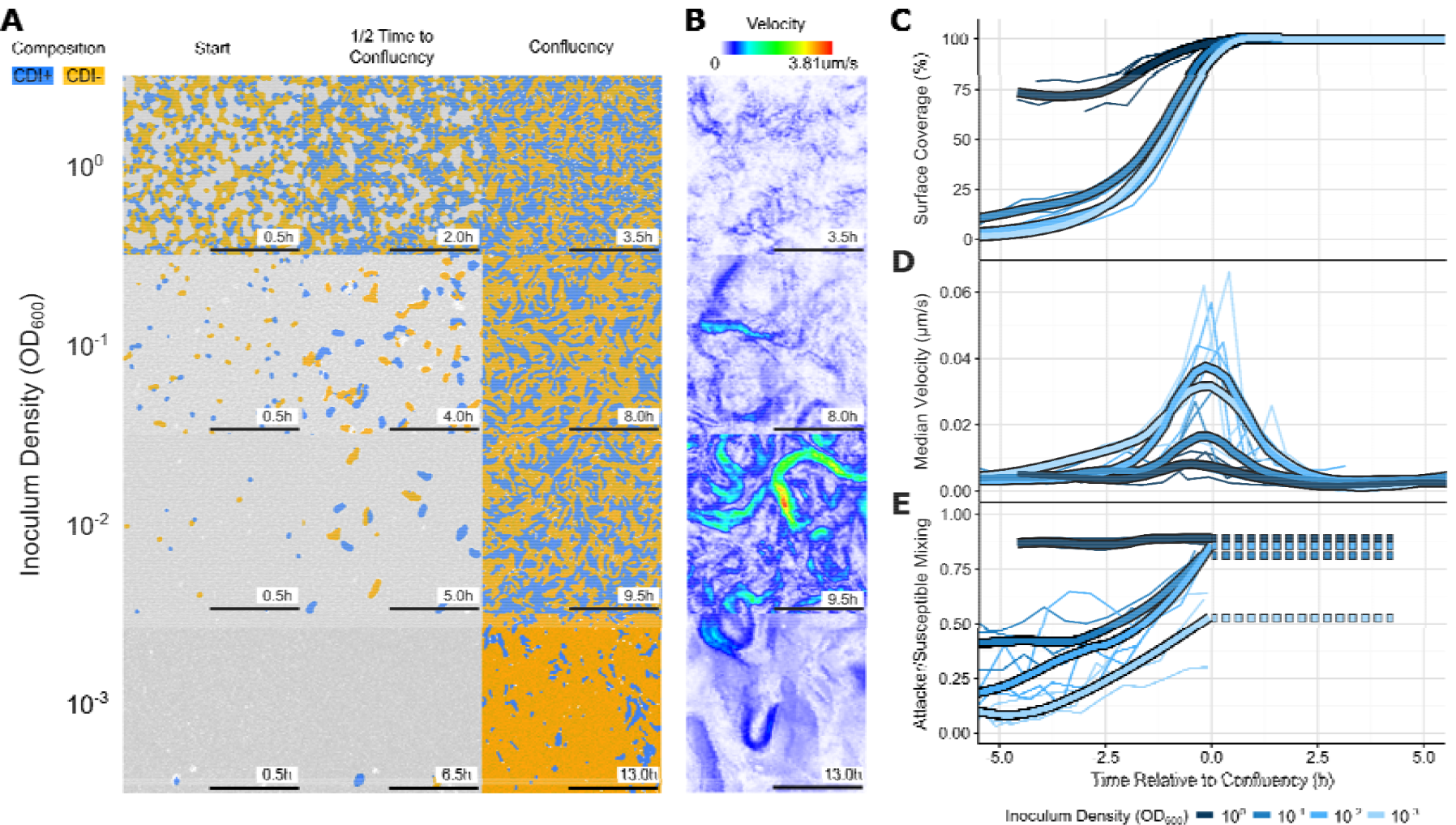
Timelapse and fluorescence microscopy show differences in dynamics of colonies inoculated at different densities. Colony competitions (1:1) between wild-type and CDI-susceptible mutants labelled with mScarlet and YFP respectively were inoculated at different initial densities (OD600) and imaged over time. Every 0.5 h after inoculation, a 1 min brightfield video was taken (0.5 frames/s) along with a single fluorescent snapshot. (**A**) Representative snapshots of colonies at the first time point (“Start”, 0.5 h), the time when the surface was completely covered by cells (“Confluency”, variable times), and halfway between the two timepoints (“1/2 Time to Confluency”, variable times) reveal increasing surface coverage with time, as well as the changing spatial distribution of wild-type attacker (blue) and CDI-susceptible (orange) strains. (**B**) The velocity fields of colonies at confluency also suggest an inverse relationship between inoculum density and cell motility. (**C-E**) To investigate these relationships further, image analysis was used to extract then plot timecourses of the percent area covered by cells (**C**), the median cell velocity magnitude (**D**), and the extent of genotypic mixing between the two strains (**E**) (for details on the image analysis see main text, methods). These revealed that the peak system velocity occurs at the time cells cover the available surface (the confluency time), up to which genotypic mixing gradually increases. To enable consistent comparisons between replicates and conditions, we centered the timecourses to this confluency time. We show individual replicates (thin lines, n=3) and LOESS-smoothed averages (thick lines) for each tested inoculum density (OD600 of 1.0, 0.1, 0.01 and 0.001). Surface coverage was fixed at 100% after reaching a peak near 99% to avoid image analysis problems associated with out-of-focus 3D colonies. We kept the genotypic mixing measurement fixed from this point for the same reason (dashed lines in E). Microscopy images have been thresholded to indicate genotypic composition.

To integrate these measurements into our theoretical framework, we used the IBM as a stepping-stone between the experimental measurements and the continuum representation of the colony; in brief, the speed, size and seeding density of cells in the experiments were matched to those of equivalent cells in the IBM, which were then converted to the continuum model parameters via our coarse-graining procedure (Fig. S2, Supplementary Note 4). We made several adjustments to the continuum model to improve the match between the experiments and the continuum model: firstly, as the genotypic mixing diffusion constant *D* and the contact switching rate *r*_*e*_ are both functions of cell velocity in the model (Fig. S2), we modified the continuum model to allow these two constants to vary as a function of time, reflecting variations in the PIV data (Fig. 4A, B). Secondly, we based the seed cell density *ρ*_0_ on images of the initial distribution of cells at each inoculum density (Fig. 4C, D). We also adjusted the model by introducing a new toxicity parameter ξthat allows us to capture an accumulating growth rate impact from each CDI intoxification event as the CDI effector used in this study is a predicted tRNAse (39) which may (reversibly (40, 41)) reduce growth rates rather than killing targets outright (14). The match between the competitive dynamics of the IBM and continuum frameworks remains strong under this model of multi-hit killing kinetics (Fig. S9). Due to the lack of experimental data, both this toxicity *ξ* and the CDI production/firing rate λ were based on biologically feasible estimates. However, our key predictions, including the non-monotonic relationship between inoculum density and competitive advantage, are robust to changes in these parameter values (Fig. S10).

**Figure 4.**
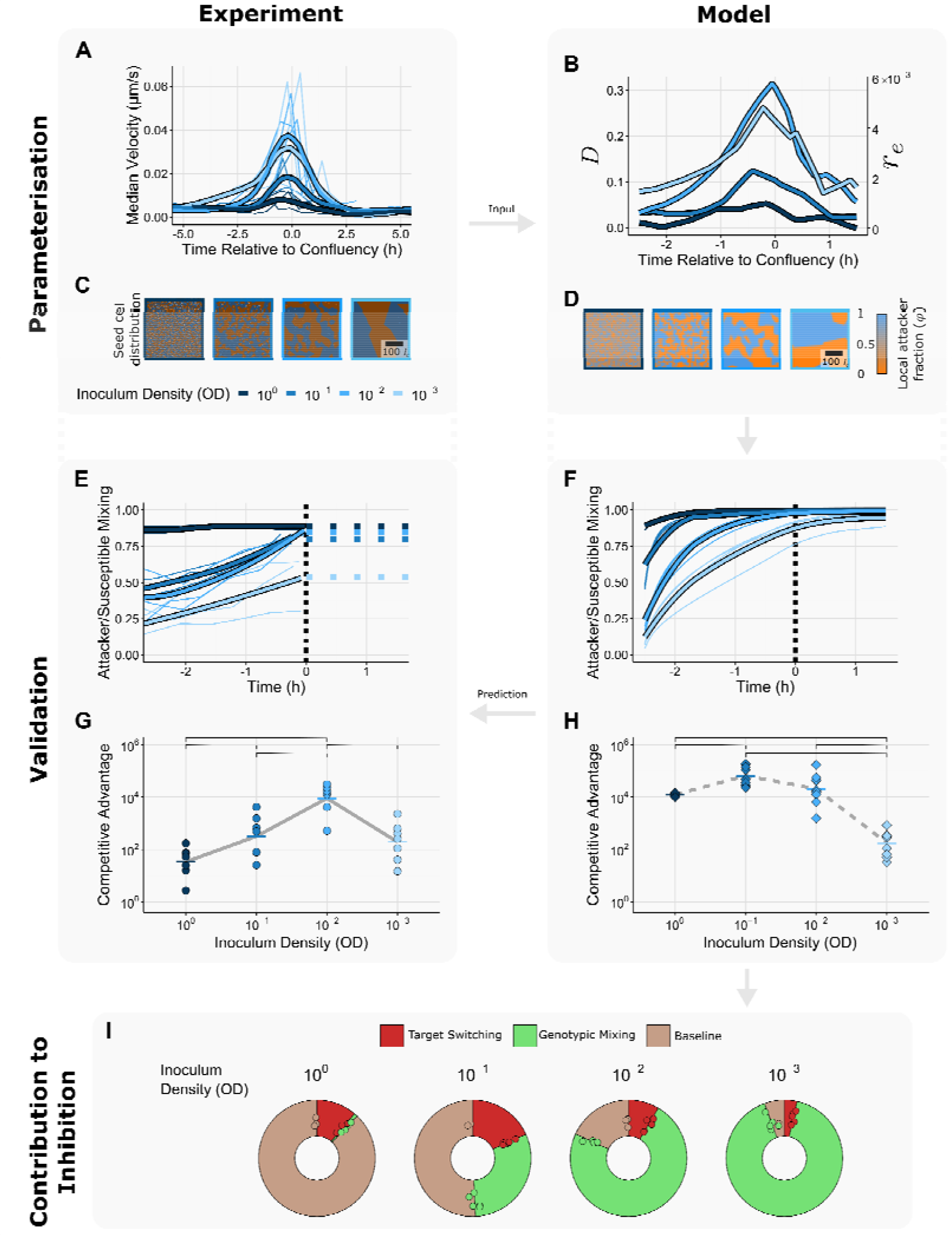
Continuum simulations matched to experiments disentangle the killing contributions of genotypic mixing and target switching. (**A-D**) We made use of the imaging data from our co-culture surface colonies to parameterise the continuum framework via the IBM representation. Velocity timecourses (**A**, reproduced from Fig. 3D) and density measurements from images of seed cells initially deposited on the agar surface (**C**, coloured regions represent corresponding Voronoi domains) were converted into the characteristic units used in the IBM, the cell width and the speed of an isolated cell with self-propulsion force. Through our coarsegraining procedure (Fig. S2), we could then transform these measurements into the continuum model parameters and (**B**), as well as starting conditions reflecting the size of clonal domains established by seed cells (**D**). By simulating the dynamics of these appropriately matched continuum models (Methods), we were able to generate predictions of the level of genotypic mixing as a function of time (**F**), as well as the final competitive advantage of the CDI+ attacker population (**H**). Comparing to the corresponding measurements from our experimental system (**E, G**, reproduced from Fig. 3D and Fig. 2C respectively), we observe that the model reproduces both the observation that the lowest inoculation density remains partially unmixed and that the relationship between inoculation density and CDI efficiency is non-monotonic. Breaking the dynamics of the continuum model into contributions from the starting conditions and the two mixing types (Methods, Fig. S3), we find that killing is mainly limited by contact switching at low inoculation densities and by genotypic mixing at high densities (**I**). F, H and I show individual datapoints and averages from n = 5 simulations initialised with different starting configurations. In G and H Lines indicate the mean of replicates (n ≥ 6). Top brackets indicate a significant difference between densities (one-sided Welch’s t-test, p < 0.05, Benjamini-Hochberg MHT corrected 0.95). See supplementary Table 2 for exact p-values.

With this framework in place, we now then used the measured velocity profiles and inoculum densities as inputs into our continuum model (Fig. 4A-D). Using these inputs, we now ran the continuum model and made predictions of the outcome of the competition under each condition.

As a first test, we compared the experimentally quantified strain intermixing to the model’s prediction (Fig. 4E, F). Importantly, as in the experiments, we observe that the lowest inoculum densities are associated with partially unmixed states at the confluency point. This result shows that our framework accurately reproduces the genotypic intermixing extent from the PIV velocity profiles and inoculum densities, indicating that our procedure for parameterising the continuum model via the IBM is effective.

Next, we calculated the predicted competitive advantage of the attacker strain across initial cell densities at the end of the experiment. Again, we find a good match between theory and data, where importantly, the model predicts the non-monotonic relationship between inoculum density and weapon efficacy (Fig. 4G, H). Our results suggest this non-monotonic relationship is due to the relative balance between motility and initial intermixing, which are affected by inoculum density in opposite ways. At high inoculum densities the two genotypes begin intermixed but experience little movement, whereas at low inoculum densities, clonal patches are large but motility is high. At intermediate densities, this trade-off between motility and initial patchiness results in an optimum that gives the maximal advantage to the attacker.

To dissect the mechanistic basis of this trade-off, we estimated the relative importance of the two motility-based mixing processes for each inoculum density (Fig. 4I). Our approach compares the baseline outcome of the continuum model when motility is not active (brown) to the outcomes when genotypic mixing (green) and target switching (red) are activated independently (Methods, Fig. S4). We observe that, similar to previous studies (5, 17, 35, 36), in the zero motility background there is a monotonic decrease in the efficacy of CDI as the inoculation density decreases and the initial clonal patches increase in size. At lower inoculum densities, the predominant mechanism by which motility improves the competitive advantage of the attacker is therefore genotypic mixing, which acts to disperse these patches. The relatively poor performance of the attacker at the lowest inoculation density arises from the failure of genotypic mixing to fully homogenise the system, as reflected in the intermixing measurements (Fig. 4E, F). By contrast, the system begins almost fully homogenised at the highest inoculation densities. Yet despite this, some sensitive cells are not initially contacted by attackers due to the stochastic process by which seed cells are deposited. Target switching therefore becomes the predominant mechanism by which motility increases the competitive advantage of the attacker, which is limited by the very low levels of motility under these conditions. In sum, our detailed analyses show that genotypic mixing and target switching can both dominate the outcome of competitions, depending on the competitive context.

## Discussion

Short-range bacterial weapons, like CDI and the T6SS, can be highly effective at killing other bacteria but suffer from severe range limitation. Here we have shown that bacteria can overcome this limitation via surface motility. When cells can move, an attacker can both better reach its targets (genotypic mixing) and avoid hitting the same targets multiple times (target switching), which solves the problem of ‘corpse-barriers’ formed by dead susceptible cells (7). In combination, these motility-driven processes greatly empower the use of contact weapons.

Similar to our findings, in Neisseria the expression of pili increases contact dependent killing by the T6SS (15). However, unlike P. *aeruginosa*, pilus expression in *Neisseria* has been found to primarily drive cell-cell aggregation. This aggregation tends to cluster pilus-expressing cells together, leading to segregation of the two genotypes when only the attacker population expresses pili and consequently reduced killing of the sensitive population (15, 42). This process, therefore, is distinct to the motility-based mechanisms we have described here, where pilus-based motility is expected to consistently result in worse outcomes for the sensitive population. Consistent with this distinction, previous work suggests that *P. aeruginosa* pili do not drive significant cell-cell adhesion in twitching cells (18). In Myxococcus *xanthus*, deletion of Type IV Pili has little effect on predation efficiency via contact-dependent mechanisms (16). However, a different mechanism, known as gliding motility, does facilitate predation in M. *xanthus*. This finding raises the possibility that the two mixing processes we describe here can be generalised across motility systems and species, as well as weapon systems. In support of this, recent work on the *P. aeruginosa* found that cell motility can improve killing via the T6SS (17).

Evolutionary game theory models predict that motility can favour the evolution of aggression. However, in these models, the process that drives aggression is the ability of motile attackers to avoid counter attacks from their targets (43, 44). We do not include the possibility of counter attacks in our study, and still find great benefits to motility, which come solely from the improved effectiveness of attacks. Adding in such benefits from avoiding counter attacks, therefore, would be likely to only further increase the benefits of motility that we have described here. Kin selection is the evolutionary process by which traits that benefit the fitness of relatives are selected for, and is strongly influenced by the genotypic similarity between nearby individuals (45). A key prediction of kin selection theory is that processes which increase interactions between individuals of different genotypes will tend to reduce cooperation and increase the potential for competition (46, 47). Our finding that genotypic mixing can greatly increase the benefits of weapon use, therefore, fits well with this prediction.

Bacteria often face strong competition for space and resources. In the face of this competition, motility can confer strong benefits by helping strains to colonise new territory (9, 48, 49). In addition, many bacteria have evolved weaponry, which enable carriers to directly inhibit and kill their competitors (1). Here we have shown that these two important mechanisms combine to generate a powerful competitive strategy that can be decisive in bacterial contests.

## Contributions

All authors were involved in the conception and writing of this study, as well as the interpretation of the theoretical and experimental results. OJM developed the IBM and continuum frameworks and performed theoretical analyses. SCB generated engineered *P. aeruginosa* strains and performed competition experiments. OJM and SCB collaboratively developed image analysis techniques.

## Acknowledgements

We would like to thank Julien Luneau, Prajwal Padmanabha, William Smith, and Jacob Palmer for feedback on an earlier version of this manuscript. K.R.F. is funded by European Research Council grant 787932 and Wellcome Trust Investigator Award 209397/Z/17/Z. O.J.M. is supported by an HFSP long-term fellowship (LT0020/2022-L).

## Data availability statement

Data are available <u>here</u>.

## Code availability statement

Code for performing IBM sweeps, continuum modelling and data analyses is available at https://github.com/Pseudomoaner/Chevauchee. The core code for the IBM is available at https://github.com/Pseudomoaner/QuasiS3R.

## Competing Interests Statement

All authors declare no competing interests.

## Materials and Methods

### Modelling

#### Individual-based model

The motion of the individual cells is governed by a pre-existing self-propelled rod (SPR) framework (18, 19) Cells are modelled as stiff chains of Yukawa segments, which repel each other via the potential *U* and self-propel at a propulsion force *F* acting along their long axis, representing type IV pili (TFP) or gliding motility. Contact intoxification occurs at a constant rate *λ* between attacker cells and contacted sensitives. Contacts are defined by drawing an ellipse two cell widths longer and wider at the same position and orientation as the attacker; any sensitive cells lying at least partially within this range are marked as contacted. Further details on IBM specification and initialisation are given in the Supplementary Methods.

#### Continuum model

Our continuum framework consists of two layers, an upper layer defining the spatial distribution of the attacker and sensitive populations, and a lower level defining the transitions of the sensitive population between different states. Tthe upper layer is a coarse-grained phase field *φ*(***r***) representing the attacker population fraction at each lattice site ***r*** (Fig. 1C). To directly compare with the IBM framework, distances are measured in cell widths; each 10x10 lattice cell therefore corresponds to a region in the IBM containing around 10 individuals. We use the semi-implicit Crank-Nicholson algorithm to simulate diffusion of this phase field with periodic boundary conditions (50), using a timestep of Δ*t* = 5.0.

The lower level is temporally finer-grained. After the distribution of attackers and sensitives has been updated, we simulate the evolution of the sensitive population at each lattice site over the next time window. To do this, we simulate transitions of the sensitive population between states indexed by a) the number of contacts made to attacker cells and b) the number of hits that have been accumulated. The transitions possible between these states are indicated in Fig. 1E. In summary, populations can gain or lose contacts with attackers one at a time, and can also accumulate individual hits. See the derivation of the explicit form of the master equation determining the behaviour of this system in Supplementary Note 1and how the continuum model parameters *r*_*e*_ and *D* were matched to the IBM data in Supplementary Note 2.

An additional parameter, *ξ* which represents the efficacy of the toxin as a percentage decrease in the growth rate of a cell due to a single CDI hit was introduced to account for the cumulative effect of multiple hits. This growth rate impact is assumed to scale linearly with the number of accumulated hits (for example, if ξ = 0.1 and we consider the sub-populations in states {5, *c*}, the total reduction in the final abundance of these sub-populations is 50 %). At the end of our continuum simulations, we have a distribution of sensitive hit/contact bins *N*_{h,c}_ at each lattice site. To convert these to a final estimate of the sensitive population size, we calculated the sum ∑ *N*_{h,c}_ (1 − ξ*h*)across all states {*h,c*} and lattice sites. The competitive index of the attacker was then calculated from this and the attacker population size as for the experiments.Comparisons between this continuum representation and our experiments required careful matching between the experimental measurements and the continuum model parameters and is described in detail in Supplementary Note 4.

### Experiments

#### Strain Construction

Deletion mutants were constructed using two-step allelic exchange using pEXG2 and conjugation with *E. coli* JKE201 (51–53). Primer sequences are listed in Supplementary Table 1. Mutants were confirmed by Sanger sequencing (Source Bioscience, Nottingham, UK). Strains were subsequently constitutively tagged with eYFP and mScarlet using pUC18-mini-Tn7-GmR (54). Approximately equal numbers of replicates were performed with each attacker/susceptible combination carrying opposite fluorescent markers. Strains are available upon request.

#### Culturing

Strains were recovered from cryo stocks onto LB 1.5% agar overnight at 30°C. Colony competitions were prepared by scraping cells off the overnight plate and resuspending cells to an initial OD600 of 1.0. Strains were mixed at defined ratios then serially diluted and 1 μL spotted on LB 1.5% agar prepared just prior. Initial culture density was determined by serial dilution followed by spot plating. Inoculum densities of OD 10^0^, 10^−1^, 10^−2^, and 10^−3^ thus correspond to approximately 2x10^6^, 2x10^5^, 2x10^4^, and 2x10^3^ cells per 1 μL droplet. Data presented are from independent colonies.

#### Quantification of Competition Outcomes

After 48 h of growth at room temperature, colonies were imaged using a Zeiss Axio Zoom V16 microscope with a Zeiss MRm camera, 0.5X PlanApo Z air objective and HXP 200C fluorescence light source. Colonies of different initial density and attacker:sensitive ratios were imaged at the same magnification, but motile and non-motile colonies were imaged at different magnifications. To make the composite images shown in the figures, the display histograms of each channel were normalized to the set of colonies sharing the same strains, meaning images can be compared within sets (i.e. motile vs non-motile) but not between. After imaging, colonies were sampled with a 10 μL pipette at both the center and edge of the colony into 0.9% saline. Samples were homogenized, serially diluted and 5 μL spotted onto LB or LB 50 μg/mL gentamycin and incubated at 30°C overnight. Colonies were counted to determine the final ratio of the two strains.

#### Calculation of Competitive Advantage

Using the initial density counted from the original inoculum cultures and the known inoculum ratios, the initial ratio of attacker:susceptible strains was determined. The final ratio was determined from the CFU counts of the serially diluted center and edge samples, with a detection limit of 10 CFU/mL used to replace zeros and prevent dividing by zero. Competitive advantage was defined as the log of the final ratio / initial ratio of attacker:susceptible cells.

#### Measurement of Cell Velocities

Bacterial cultures were prepared as previously and 1 μL of a 1:1 mixture of wild-type and CDI susceptible pipetted onto the center of a glass bottom petri dish (MatTek, Goethenberg, Sweden) containing 10 mL LB 1.5% agar. As soon as the droplet was dry, a lid with a coverslip was attached, sealed with parafilm and the sample inverted onto the microscope (Zeiss Axio Observer, 20X Air objective, Colibri LED lightsource). Every 0.5 h, one fluorescent image was taken including the brightfield, then a 1 minute timelapse of just the brightfield, with images taken every 2s. Before taking the first image at each timepoint, the focus was adjusted to account for the vertical drift of the sample over time using a combination of definite focus (Zeiss Definite Focus 2) and manual correction.

#### Observation of contact exchange switching

To observe individual cells exchanging contacts due to motility, surface colonies were prepared as above for measuring cell velocity, but were only imaged just prior to confluency being reached. These colonies were imaged using a 50X air objective with the coverslip removed. Fluorescent images were taken every 30 s, while brightfield images were taken approximately every 1 s for 5 minutes. The focus was manually adjusted for the entire length of the timelapse to account for drift. To track individual cells, the brightfield timelapses were initially segmented using FAST (55) and manually corrected. Contacts of individual cells were assessed manually.

### Data analysis

#### Image Pre-Processing

To prepare the 1 minute timelapses for particle image velocimetry (PIV), they were divided by a gaussian smoothed (5px) median projection of the entire timelapse. Fluorescent images were corrected for uneven illumination based on a computed flat field using BaSic (56).

#### Particle Image Velocimetry

PIVlab (37, 38) was used to estimate the velocity of motile cells over time. Briefly, pre-processed images were loaded into PIVlab, and velocity field profiles were calculated for each pair of subsequent frames from the 1 minute brightfield timelapses. Exact PIVlab settings are specified in the scripts linked to in the code availability statement. The signal in cell-free regions was excluded using based on low inter-frame correlation scores. Velocity vectors were averaged over the entire timelapse to isolate the net cell motion from the rapid back-and-forth motion of twitching cells, then the magnitudes of these average vectors were determined. Median velocity magnitude was used as a summary statistic that represents the amount of motility occurring in a colony at each timepoint.

#### Fluorescent Image Processing

Fluorescent images were used to determine agar surface coverage mixing of attacker and sensitive cells. Using adaptively adjusted thresholds to smooth our variations from instabilities in the focal plane, the background was segmented out using a texture-based metric of the brightfield channel yielding surface coverage, and the remaining pixels assigned as either attacker or sensitive using the log-transformed pixel-wise ratio of eYFP/mScarlet intensity. Mixing was then calculated by overlaying a lattice of boxes and calculating the attacker fraction in each box that was at least 5% occupied using the formula 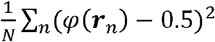. This variance reaches a maximum of 0.25 when the strains are fully segregated and a minimum approaching 0 when strains are fully intermixed. To intuitively show these data to 0-1 by subtracting 0.25 then dividing by -0.25, making 0 completely unmixed and 1 completely mixed. The intermixing measurements for the continuum model were calculated similarly, using the attacker fraction *φ*(***r***) output by the model directly.

## Notes

https://github.com/Pseudomoaner/Chevauchee

https://github.com/Pseudomoaner/QuasiS3R

https://figshare.com/articles/dataset/Data_for_Booth_Meacock_Foster_2023/23822169

### Competing Interest Statement

The authors have declared no competing interest.

