## Supplementary information for "Cell motility greatly empowers bacterial contact weapons"

Sean C. Booth, Oliver J. Meacock, Kevin R. Foster

### Supplementary Methods

#### Individual-Based Model

##### Intoxification

Although some weapon systems (such as the T6SS) are spent as soon as they are fired regardless of whether they hit a target, others (such as CDI systems) are likely to be stored on the cell surface in an untriggered state and then get distributed among contacted targets. To test whether this difference in toxin usage dynamics has a substantial impact on intoxification rates, we performed separate simulations in which $\lambda$ was either kept constant or else was scaled by the instantaneous number of contacts to the attacker. We found that intoxification under such dynamic contact scaling was almost identical to what we would predict from reducing the constant intoxification rate by the average number of cell contacts (Fig. S1). As this is a subset of our constant-rate simulations, we therefore only consider the constant firing rate form of the simulations elsewhere in this manuscript.

##### Equations of motion

Taking the instantaneous position of a rod $\alpha$ as $\boldsymbol{r}_\alpha$, its orientation as $\theta_\alpha$, the unit vector denoting this orientation as $\hat{\boldsymbol{u}}_\alpha$ and the sum of the potentials between $\alpha$ and all other rods as $U_\alpha$, the equations of motion for each rod in the IBM are:

$$\boldsymbol{f}_T \cdot \frac{\partial \boldsymbol{r}_\alpha}{\partial t} = -\frac{\partial U_\alpha}{\partial \boldsymbol{r}_\alpha} + F\hat{\boldsymbol{u}}_\alpha, \qquad \text{(S1a)}$$

$$f_\theta \frac{\partial \theta_\alpha}{\partial t} = -\frac{\partial U_\alpha}{\partial \theta_\alpha}. \qquad \text{(S1b)}$$

Here $\boldsymbol{f}_T$ is the translational friction tensor, which allows simulation of non-isotropic friction along the long and short axes of the cell, and $f_\theta$ is the rotational friction constant. $\boldsymbol{f}_T$ and $f_\theta$ are themselves functions of the rod aspect ratio and the Stokesian friction coefficient, $f_0$, which we define using the formulation presented in [1].

##### Simulation specifications

Motility is simulated by using the midpoint method to numerically integrate Eqs. 3a and 3b. We use a fixed timestep of $\Delta t = 0.1$, which produces numerically stable results for all simulation conditions discussed in this paper. The system density, defined as the total cell area divided by the area of the simulation domain, was 0.35. Following the approach of [2], we non-dimensionalise our simulations by a) setting the characteristic length scale of the system to be a single cell width, and b) setting $f_0 = 1$. Simulations were performed in a 200x200 simulation domain with doubly periodic boundary conditions. Other parameters are specified in the relevant figures.

As calculating contacts is a computationally intense process, we performed contact detection and hit application on only a subset of frames ($\Delta t = 1.0$). Nonetheless, this was sufficiently fine-grained to accurately capture hit accumulation.

##### Initialisation

Model initialisation proceeds in two phases. To begin with, we initialise all cells on a lattice, each individual being oriented randomly left or right. We then allow the system configuration to reach a statistical steady-state by turning on cell movement and running the simulation dynamics until statistical measures of the system (specifically, the distribution of cell orientations) stabilise.

In the second phase, we impose genotypic patchiness on this equilibrated cell field. We simulate different starting inoculation densities by choosing random points within the simulation domain with density $\rho_0$. Each point is assigned either to the attacker or sensitive population, with the fraction of attacker points being specified by the user. These points are intended to represent seed cells that are laid down on the surface at the beginning of the experiment. In experimental systems, these grow into clonal patches that are well approximated by a Voronoi tessellation over the seed points [3]. To reproduce this effect in the IBM, we construct a Voronoi tessellation based on these seeding points and assign all the cells within the equilibrated field that fall within a particular patch to that patch's population. Beyond this point, motility and intoxification are simulated concurrently.

#### Experiments

##### Fluorescent Image Processing

Fluorescent images were used to determine how much of the agar surface was covered in cells at each timepoint and how mixed the attacker and susceptible strains were. Cells were segmented from the background by using a texture-based metric on the brightfield channel to distinguish cell-containing and cell-free regions. This segmentation was then further broken down into attacker and sensitive segmentations by using it as a mask on the two sets of fluorescence images to obtain the eYFP and mScarlet intensities of pixels within cells. We then took the pixel-wise ratio of these intensities and log-transformed the result, yielding a histogram with two peaks corresponding to the eYFP and mScarlet-labelled populations. These peaks were divided by finding the location of the local minimum between them, allowing us to automatically assign pixels to the two populations. After an initial pass through all the data to obtain estimates of the packing fractions and population fractions, both the texture and fluorescence thresholds were adaptively adjusted to smooth out variations of these metrics caused by instabilities in the focal plane.

Surface coverage was calculated as the ratio of the area of the brightfield segmentation to the total image area. Strain mixing was calculated from the fluorescence images by overlaying a lattice of boxes equivalent in granularity to the coarse-grained lattice of the continuum model (i.e. each box was 10 cell widths x 10 cell widths), then determining the attacker fraction $\varphi(\boldsymbol{r})$ in each of the $N$ boxes and calculating the variance across the whole field of view. Assuming a global 50:50 ratio of the two populations (the inoculation ratio), this calculation is expressed as $\frac{1}{N}\sum_n(\varphi(\boldsymbol{r}_n) - 0.5)^2$, where the summation occurs over all included lattice sites $n$. Boxes less than 5% occupied – as estimated from the brightfield segmentation – were removed for this calculation, allowing us to discount the large regions of empty space in the lowest inoculation density colonies at early timepoints (Fig. 3A). This variance reaches a theoretical maximum of 0.25 when cells are fully segregated and a theoretical minimum close to 0 when genotypes are fully intermixed, so to intuitively show these data (Figs. 3E, 4E, S8C) they were normalized to a range of 0-1 by subtracting 0.25 then dividing by -0.25, making 0 completely unmixed and 1 completely mixed. The intermixing measurements for the continuum model (Fig. 4F) were calculated similarly, using the attacker fraction $\varphi(\boldsymbol{r})$ output by the model directly.

##### RNA Extraction

RNA was extracted using a RiboPure RNA purification kit (Bacteria) (ThermoFisher). Colonies were inoculated as normal (from 30C overnight streaked cells were scraped off the plate and diluted in 0.9% saline to OD1, 1uL spotted on fresh LB 1.5% agar poured then dried in laminar flow hood for 15 minutes). After 24h of growth at RT, colonies were scraped off directly into 350uL RNAwiz, then disrupted to homogeneity by pipetting up and down. The kit instructions were then followed: samples were added to 250uL pre-aliquoted zirconia beads then bead beat using a Fastprep-24 5g (MP Biomedicals) beating for 40 s at 6 m/s twice with a 5 minute wait (on ice) between beatings. An extraction blank with no input sample was also processed. Samples were eluted in 50uL pre-warmed elution solution which was then run through the membrane again for a second elution. The optional DNAse treatment was then performed and RNA concentration and purity was determined using a Nanodrop 1000 (Thermo Scientific). RNA was stored at -20C until further analysis.

#### q-RT-PCR

A Luna Universal One-step RT-q-PCR kit (NEB) and Stratagene Mx3005P qPCR thermocycler (Agilent) were used to quantify expression of the CDI toxin gene. The kit protocol was followed using 2uL of input RNA from each sample and either CDI primers (14) or acpP (55). With each primer set, three technical replicates of each test sample were prepared, as well as a no-template control, positive control and single technical replicate of the extraction blank. Thermocycling conditions were as specified by the manufacturer and the melting curve was from 60 - 90C. Cycle thresholds were calculated using the MXPro software v4.10 (Amplification-based threshold, adaptive baseline, moving average: 3), using the SYBR signal normalized to ROX. CDI expression is reported as the cycle threshold (Ct) minus the Ct of acpP (ΔCt). Melting curves were plotted in R using ggplot2 (56) with a geom_smooth span setting of 0.1.

### Supplementary Notes

#### Supplementary Note 1 | Derivation of continuum model master equation

As discussed in the main text and the Methods, we simulate transitions of populations at a given lattice site $\boldsymbol{r}$ between different states (labelled by the number of accumulated hits $h$ and number of attacker contacts $c$) as a Markovian process. We can construct a master equation for this system by determining the rates of these transitions. Using the notation $\omega(\{1\} \longrightarrow \{2\})$ to denote the rate of the transition from state $\{1\}$ to state $\{2\}$, examination of Fig. 1E and the generic master equation (Eq. 2) suggests an equation of the form

$$\begin{aligned}\frac{dN_{\{h,c\}}}{dt} = \; & N_{\{h-1,c\}}\omega(\{h-1,c\} \longrightarrow \{h,c\}) - N_{\{h,c\}}\omega(\{h,c\} \longrightarrow \{h+1,c\}) \\ & + N_{\{h,c-1\}}\,\omega(\{h,c-1\} \longrightarrow \{h,c\}) - N_{\{h,c\}}\omega(\{h,c\} \longrightarrow \{h,c-1\}) \\ & + N_{\{h,c+1\}}\,\omega(\{h,c+1\} \longrightarrow \{h,c\}) - N_{\{h,c\}}\omega(\{h,c\} \longrightarrow \{h,c+1\}) \end{aligned} \qquad \text{(S2)}$$

This is simply the rate of each step into the state $\{h, c\}$ multiplied by the size of the population acting as the source for that transition, minus the rate of each step out of $\{h, c\}$ multiplied by the size of the population in that state ($N_{\{h,c\}}$).

The hit accumulation rate is set by the parameter $\lambda$. This is scaled by $c$, the number of attackers currently contacting this population of sensitives. Thus,

$$\omega(\{h,c\} \longrightarrow \{h+1,c\}) = \omega(\{h-1,c\} \longrightarrow \{h,c\}) = c\lambda. \qquad \text{(S3)}$$

To define the contact exchange rates, we begin by noting that individual contacts are broken and formed at the rate $r_e/C$, where $C$ is the total number of contacts made by each sensitive cell (in general in this manuscript $C = 5$, Fig. S2A). There are then four possible scenarios for a given contact exchange:

- A sensitive-sensitive contact is broken and a sensitive-attacker contact is made ($\{h,c\} \longrightarrow \{h,c+1\}$). The probability of this is the current fraction of sensitive-sensitive contacts multiplied by the attacker fraction at this lattice site, which is $\left(1-\frac{c}{C}\right)\varphi(\boldsymbol{r})$.
- A sensitive-sensitive contact is broken and a sensitive-sensitive contact is made ($\{h,c\} \longrightarrow \{h,c\}$). The probability of this is $\left(1-\frac{c}{C}\right)(1-\varphi(\boldsymbol{r}))$.
- A sensitive-attacker contact is broken and a sensitive-sensitive contact is made ($\{h,c\} \longrightarrow \{h,c-1\}$). The probability of this is $\frac{c}{C}(1-\varphi(\boldsymbol{r}))$.
- A sensitive-attacker contact is broken and a sensitive-attacker contact is made ($\{h,c\} \longrightarrow \{h,c\}$). The probability of this is $\frac{c}{C}\varphi(\boldsymbol{r})$.

As $\{h,c\} \longrightarrow \{h,c\}$ transitions are 'silent' (result in no change in state), we only need to consider the first and third cases. Combining with the target switching rates and accounting for $C$ contact sites, these yield

$$\omega(\{h,c\} \longrightarrow \{h,c+1\}) = r_e\left(1-\frac{c}{C}\right)\varphi(\boldsymbol{r}), \qquad \text{(S4a)}$$

and

$$\omega(\{h,c\} \longrightarrow \{h,c-1\}) = r_e\frac{c}{C}(1-\varphi(\boldsymbol{r})). \qquad \text{(S4b)}$$

We can plug these terms into our master equation and perform some algebraic manipulations to obtain

$$\frac{dN_{\{h,c\}}}{dt} = c\lambda\big(N_{\{h-1,c\}} - N_{\{h,c\}}\big)$$
$$+\frac{r_e}{C}\Big((c+1)\,(1-\varphi)N_{\{h,c+1\}} + (C-c+1)\varphi N_{\{h,c-1\}} - (c(1-\varphi) + (C-c)\varphi)N_{\{h,c\}}\Big). \qquad \text{(S5)}$$

In practice, to avoid keeping track of a potentially unlimited collection of separate populations, we set a cap on the maximum number of hits that are explicitly tallied. We set the rate $\omega(\{H,c\} \longrightarrow \{H+1,c\}) = 0$, meaning in effect that the set of populations $\{H,c\}$ contains all sensitives with $H$ *or more* hits. The value of $H$ depends on the potency of the toxin $\xi$, such that it corresponds to the number of hits necessary to set a cells growth rate to zero. These ODEs are numerically integrated using Matlab's ode45 solver at each lattice site.

#### Supplementary Note 2 | Parameterization of continuum model from IBM data

Direct comparison between the IBM and the continuum frameworks requires that we obtain estimates of the key parameters $r_e$ and $D$, which define the rates of target switching and genotypic mixing respectively. We can estimate these parameters directly from cell tracks generated by the IBM.

$r_e$ can be measured simply as the average rate of new contact formation (Fig. S2A). Because contacts can break and reform repeatedly when cells are at the edge of the contact region, we require that contacts remain stable for at least 5 time units to be included in the final count. Neglecting to do this would result in a substantial overestimate of the contact rate.

To estimate $D$, we calculate the Root Mean Squared Displacement (RMSD) $r(\tau)$ of simulated cells along their trajectories. Explicitly,

$$r(\tau) = \langle|\boldsymbol{x}(\tau + t) - \boldsymbol{x}(t)|^2\rangle^{\frac{1}{2}}, \tag{S6}$$

where $\boldsymbol{x}(t)$ denotes a cell's position vector at time $t$, $\tau$ is some query lag time and the angular brackets $\langle\cdot\rangle$ denote the ensemble average over all cells and $t$. Microorganisms typically have two parts to their RMSD curve; at small lag times $\tau$, cells move ballistically in straight lines, while at longer lag times, reorientations of cells due to Brownian motion and interactions with other individuals leads to patterns of net motion resembling diffusion [4]. In two dimensions, this leads to the relationship

$$r(\tau) \approx 2\sqrt{D\tau} \tag{S7}$$

at long timescales. On log-transformed axes, this is represented as a curve with a slope of ½ at long lag times.

We observe this in our own IBM data (Fig. S2B), with the gradient of the RMSD curve declining at longer lag times. However, due to the finite size of the simulation domain, some of the high-force simulations show sub-diffusive behaviour at the very longest lag times. To address this, we find $D$ by using the tangent to the RMSD curve where its log-transformed gradient is exactly ½, revealing an approximately linear relationship between $\bar{v}$ and $D$ (Fig. S2B, inset).

It is interesting to note that we observe a linear scaling between these two variables – typically, we would expect $D = \bar{v}^2 t_c/2$ for free-living planktonic organisms in two dimensions, where $t_c$ is the timescale at which the transition from ballistic to diffusive motion occurs [4]. The reason for this apparent discrepancy appears to be due to the highly crowded conditions of the IBM. In contrast to planktonic cells, reorientations of cells in this system occur due to pushing by neighbouring cells, rather than Brownian jostling from the surrounding fluid. As the rate of this pushing by neighbours will depend upon their velocity, we would expect that $t_c$ is not constant but is itself inversely proportional to $\bar{v}$, leading to the observed scaling.

#### Supplementary Note 3 | Analysis of dependence of attack efficiency on velocity and firing rate

In the main text, we focus on the influence of cell velocity and starting system patchiness on intoxification efficiency, as these are parameters that can be measured and modified in our experimental system. However, there are other factors which can influence the efficiency of intoxification. In this note, we derive a relationship that illustrates one such factor, showing that the firing rate of the contact-dependent weapon determines how quickly intoxification saturates as a function of cell velocity.

We begin by assuming that we are under homogeneous conditions, with no patchiness. Under these conditions contact exchange becomes the only velocity-limited process. We further assume that $\sum_h \sum_c N_{\{h,c\}} \gg \varphi$ (*i.e.,* that there are many more sensitive cells than attackers), which corresponds to invasion of a sensitive population by a small population of attackers. This implies that there will be almost no sensitive cells contacted by more than one attacker, meaning we need to keep track of two contact bins (sensitives contacted by a single attacker or not contacted by attacker). Finally, we assume that a single hit is sufficient to inactivate a target sensitive.

A schematic of this simplified formulation is shown in Fig. S3A. This scheme is highly suggestive of an enzymatic catalysis, and indeed under the assumptions listed above we can make use of the Briggs-Haldane formalism to write

$$\frac{d\sum_c N_{\{0,c\}}}{dt} \approx \lambda\varphi C \frac{\sum_c N_{\{0,c\}}}{K_M + \sum_c N_{\{0,c\}}} = \lambda\varphi C \frac{\alpha_e \bar{v} \sum_c N_{\{0,c\}}}{\alpha_e \bar{v}\left(1 + \sum_c N_{\{0,c\}}\right) + \lambda}. \qquad \text{(S8)}$$

where the notation $\sum_c N_{0,c}$ indicates the total proportion of unhit sensitive cells. The closed-form solution to this equation is transcendental, and not amenable to easy interpretation [5]. However, we can gain considerable insight by considering the intoxification rate at $t = 0$, which will remain approximately constant for as long as the proportion of unintoxified sensitive cells remains (approximately) unchanged. Then, under the assumptions listed above, we have:

$$\frac{d\sum_c N_{\{0,c\}}}{dt} \approx \lambda\varphi C \frac{\alpha_e \bar{v}}{2\alpha_e \bar{v} + \lambda}. \qquad \text{(S9)}$$

This expression is plotted in Fig. S3B, along with values of $\frac{d\sum_c N_{\{0,c\}}}{dt}$ estimated from IBM simulations run with low attacker fractions and homogeneous starting conditions.

Both the IBM simulations and this analytical approximation agree on several points. Firstly, and most basically, increasing velocity is predicted to always improve intoxification efficiency. However, this intoxification rate is a saturating function of system activity, with larger increases in cell velocity providing increasingly small benefits to the attacker. Mechanistically, this saturation occurs because in the high-velocity limit contacts are exchanged before attackers have an opportunity to intoxify the target. Attackers are therefore guaranteed to be in contact with an unhit sensitive cell during their rare firing events, and the overall rate of intoxification is determined by this firing rate. Finally, the half-maximal intoxification rate is obtained at $\bar{v}_{\frac{1}{2}} = \frac{\lambda}{2\alpha_e}$. It is therefore predicted that attackers can overcome this velocity-dependent saturation effect by simultaneously increasing their firing rate.

#### Supplementary Note 4 | Matching experiments to the continuum framework

In order to compare our experimental data with the continuum model, we first needed to appropriately parameterize the continuum model via the IBM. To do this, we needed to match the physical units used to measure experimental quantities to those used in the IBM and thence to those of the continuum model. We chose 1 cell width (or $l_c$) as the characteristic lengthscale of the models to use as the base unit for spatial measurements, corresponding to 0.8 µm in our images. As the continuum model lacks a characteristic timescale, we arbitrarily chose 1 second as our units of time, meaning velocities were measured in units of $l_c$ $s^{-1}$ (choice of another time unit as the basis of the simulations, *e.g.* 1 minute, merely rescales the dynamics of the model and does not change its final output). Finally, to match the initial seeding densities in the experiments and the models, we manually counted the number of seed cells present per unit area for different inoculum ODs. As expected, we found that the turbidity of the inoculum was linearly proportional to the seeding density, with a proportionality constant of 0.1 cells $l_c^{-2}$ $OD_{600}^{-1}$. Together, these considerations were sufficient to match the three parameters of the continuum model $\rho_0$, $r_e$ and $D$ to our experiments (the latter two being velocity-dependent, Figs. 4B, S2).

Both the toxin efficacy parameter $\xi$ and the hit rate parameter $\lambda$ were difficult to estimate from existing studies; Bottery et al. [6] report a modest reduction in growth rate for sensitive *E. coli* cells growing adjacent to CDI+ cells (~30%), but as no method currently exists for quantifying CDI firing rate it is difficult to estimate the potency of a single CDI hit from this data. We also note that equivalent measurements are not currently available for the organism we use here, *P. aeruginosa*. We therefore performed a parameter sweep over these two quantities to ensure that our main findings were robust to variations in them (Fig. S10). Although the parameter choices presented in the main text ($\lambda = 0.005$ $s^{-1}$ and $\xi = 0.02$) give the closest match to our experimental results in absolute terms, crucially we find that the non-monotonic relationship between inoculation density and the competitive index of the attacker is preserved across most of the parameter space.

Finally, we note that we restrict the simulation time to a window around the peak in motility, from 2.5 hours before the confluency time to 1.5 hours after this timepoint. This window is symmetric about the peak in motility, which occurs around 0.5 hours before confluency (Fig. 4A). We perform this windowing to minimise the discrepancy between the varying density of the experiments and the fixed density of the continuum system.

##### Assumptions of matching procedure

We rely on several simplifications to perform this comparison between the experimental data and our continuum framework. In the following, we explicitly state these assumptions and provide justifications for them.

- The continuum formulation treats the density of the system as constant, rather than a quantity that increases over time as we observe in our data (Fig. 3C). Our reasoning behind this assumption is based on the expectation that the primary impact of CDI will occur during the window that cells are motile; both before and after this time, structures (both corpse barriers and clonal patches) will be 'frozen in', and contact-based weapons will be rendered ineffective. This is supported by the observation that motility is the dominant factor determining the outcome of the competition across inoculation densities and ratios (Fig. S6B). Motility is in turn restricted to a limited time window (Fig. 3D) during which only around 2-3 cell divisions take place, as estimated by the change in surface coverage. Thus, restriction of the simulation of the system within this sub-window is expected to provide a reasonable approximation of the outcome of the whole experiment. While the observed ~4-fold increase in the density of the experimental colonies over this window is certainly substantial, it is small relative to the effects on competitive outcome – up to a 10,000-fold difference in final abundances of the sensitive population in the motile vs. non-motile cases - that we observe.

- We also assume that the genotypic structure of the system at confluency is well-approximated by the Voronoi-based approach described above, even in the presence of motility. We base this on the observation that even in the presence of motility, clonal patches remain largely distinct until a couple of hours before confluency (*e.g.* Fig. 2D). This is reflected in our measurements of genotypic mixing, which increase slowly up to 2.5 hours before confluency and more rapidly after this point (Fig. 3E). Patches therefore begin to substantially mix at around the same time motility begins to be consequential, which is the time at which we initialise our continuum simulations.
- Finally, we assume that the slowdown in growth rate mediated by contact-dependent hits only impacts a cell for a single generation, with later generations returning to the baseline growth rate. This assumption lies behind our assumption that the final abundances of different sub-populations can be rescaled by $(1 - \xi h)$, rather than, for example, $2^{-n\xi h}$, which would be the expected expression if all offspring inherited the growth rate impact of the toxin from the originally hit cell (in this expression, $n$ is the number of divisions between the intoxification time and the final timepoint). We justify this by noting that cells subjected to several CDI systems, including tRNAase-based systems such as those we use here, revert to their original growth rates within a few hours of removal of inhibition [7,8]. In reality, the true impact of the CDI toxin most likely lies between these two extremes of single-generation vs. indefinite toxicity, however we note that our assumption is conservative in the sense that multi-generation toxic effects would be expected to increase the overall impact of CDI.

### Supplementary Tables

#### Supplementary Table 1 | Primers used for construction of Δ*pilU* strains and q-RT-PCR

| Primer | Use | Sequence |
|---|---|---|
| pilU_Up_F | *pilU* upstream region | CAAGCTTCTGCAGGTCGACTCTAGAGGATCgcagaccctgatcaagaagatcg |
| pilU_Up_R | *pilU* upstream region | gcctactgaagacggttcagcggaagcgccattccatgatgttctcgctcactc |
| pilU_Down_F | *pilU* downstream region | gcccctgagtgagcgagaacatcatggaatggcgcttccgctgaacc |
| pilU_Down_R | *pilU* downstream region | ACCCGTGGAAATTAATTAAGGTACCGAATTtcggcgtggccttctatatcc |
| acp_F | *acpP* qPCR | ACTCGGCGTGAAGGAAGAAG |
| acp_R | *acpP* qPCR | CGACGGTGTCAAGGGAGT |
| CDI1_F | CDI 1 qPCR | cgcgatgaaggcaacctgc |
| CDI1_R | CDI 1 qPCR | ccgtggacgttcaactcg |

### Supplementary Figures

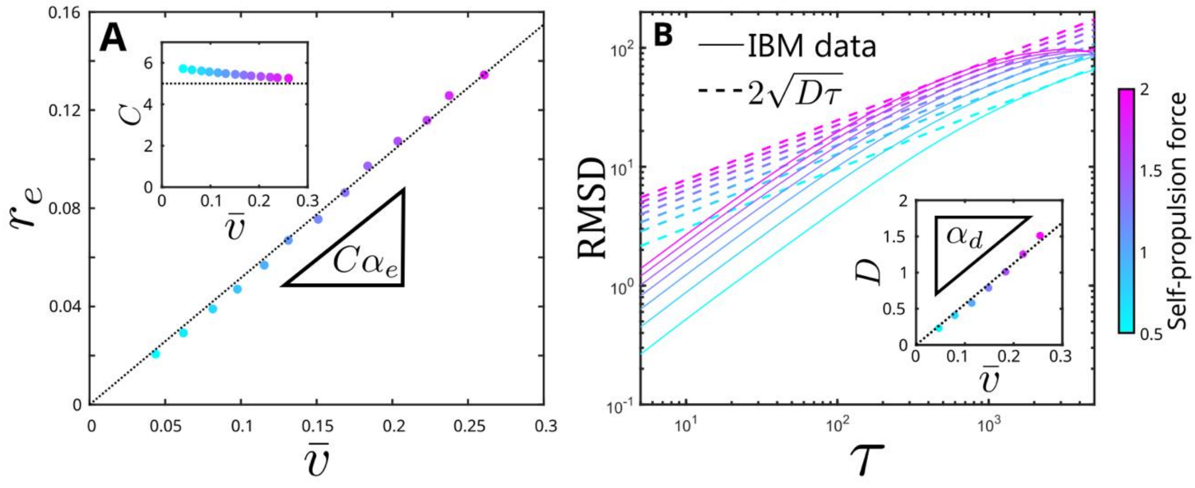

**Figure S1 | Continuum model parameters can be extracted from the IBM.** To find the relationships between average cell speed $\bar{v}$ and the target switching rate $r_e$ and diffusion constant $D$ associated with genotypic mixing, we extracted relevant statistics from self-propelled rod simulations run with different self-propulsion forces. (**A**) We directly measured the rate at which cells made contact with new individuals, yielding the per-cell target switching rate $r_e$. This is the product of the per-contact contact switching rate and the total number of cell contacts $C$, which is weakly dependent on system activity (inset), probably due to activity-dependent density fluctuations [9]. We assume $C = 5$ (dotted line) throughout this manuscript – the closest integer value under most conditions – although we obtain substantially similar results for $C = 6$. (**B**) We further measured the diffusion constant of simulated cells by calculating their Root Mean Squared Displacement (RMSD). We estimate the effective diffusion constant $D$ of cells at these long timescales by finding the tangent to the log-transformed RMSD data with a slope of 1/2 (dashed lines). In both **A** and **B**, we found a simple proportional relationship between $\bar{v}$ and the statistic in question. We fit the proportionality constants $\alpha_e$ and $\alpha_d$ to these, allowing us to objectively parameterise the connection between our IBM and continuum frameworks (Fig. 1).

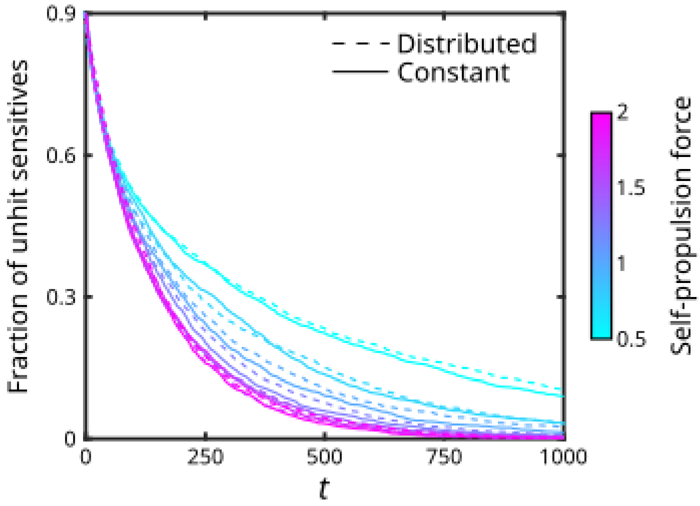

**Figure S2 | Different microscopic firing mechanisms result in equivalent macroscopic killing dynamics.** To investigate the importance of the mechanistic basis of different types of contact-dependent weapons for determining killing dynamics (Methods), we performed IBM simulations with different intoxification processes. In the 'Distributed' simulations, we assumed that surface-based toxins were produced stochastically at a rate $\lambda$ and distributed equally between all cells currently contacted, representing a delivery mechanism such as CDI where the toxin can remain active on the cell surface until it is taken up by a neighbouring cell. In the 'Constant' simulations, all currently contacted cells were stochastically intoxified at a fixed rate $\lambda$, representing a toxin delivery mechanism such as the T6SS where delivery is only possible at the time of firing. We find that the overall intoxification dynamics are equivalent between the two mechanisms, provided the firing rate $\lambda$ was rescaled by the average number of cell contacts $C = 5$ ($\lambda = 0.1$, Distributed, $\lambda = 0.02$, Constant). Simulations were initialised under homogeneous (non-patchy) starting conditions, with an initial fraction of attackers of 0.1.

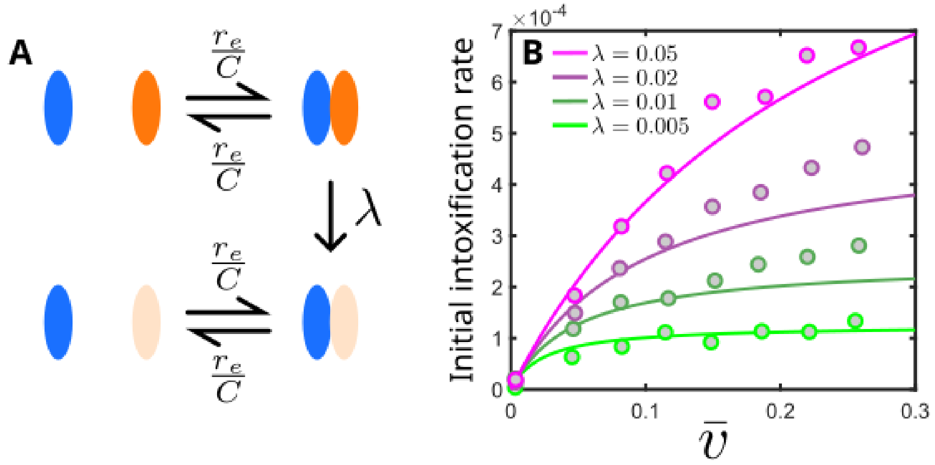

**Figure S3 | Intoxification efficiency is a saturating function of system velocity under homogeneous starting conditions.** (**A**) Under homogeneous (non-patchy) conditions, contact exchange is the sole velocity-dependent process limiting intoxification efficiency. We can model the resulting intoxification dynamics using an enzyme-like reaction scheme, with the per-cell contact exchange rate $r_{e=}C\alpha_e\bar{v}$, number of cell contacts $C$ and weapon firing rate $\lambda$ setting the various rates in the system. (**B**) From this, we can predict the initial intoxification rate of a small invading population of attackers as a function of the average cell velocity $\bar{v}$ and weapon firing rate $\lambda$ (Supplementary Notes). Solid lines are predictions from the analytical model, points are IBM outputs.

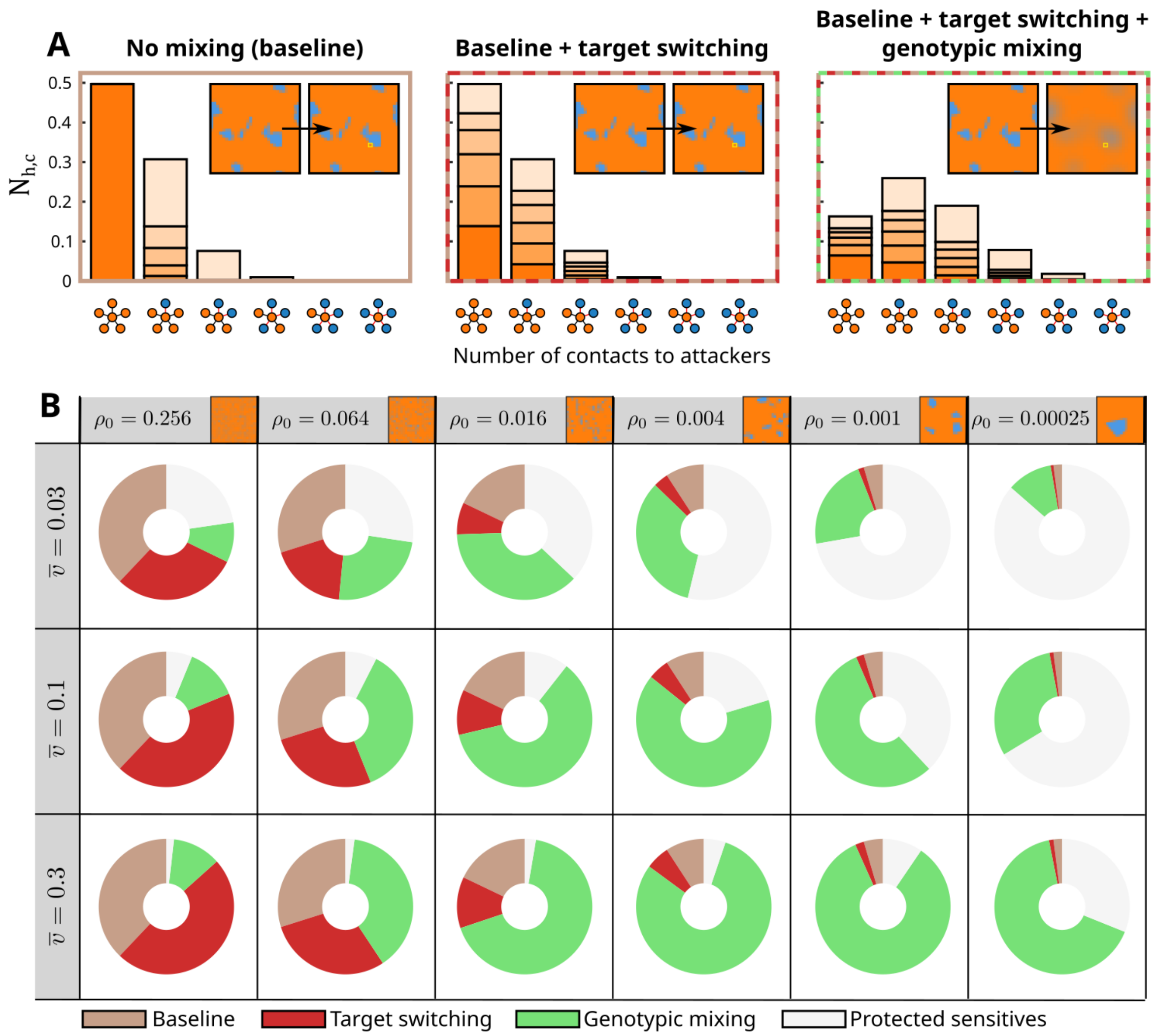

**Figure S4 | The continuum framework allows us to disentangle contributions from the two forms of mixing.** (**A**) From equivalent starting conditions, continuum simulations can be run with no mixing (left), only contact switching active (middle) or with both contact switching and genotypic mixing active (right). (**B**) By comparing the number of cells intoxified under each of these conditions, we can split the intoxification dynamics under a given set of conditions into a baseline component guaranteed by the starting conditions even in the absence of mixing (brown), a component mediated by contact switching (red) and a component mediated by genotypic mixing (green). In addition, a portion of the sensitive population remains unaffected even when both forms of mixing are active (light grey). Here we show the variable contributions of the two mixing processes when varying motility and patchiness. In (**A**), the contact distributions shown are taken at *t* = 1000 of a simulation run with the specified mixing types active. The lattice site sampled to generate these contact distributions is highlighted with a yellow square in the insets. In all cases, simulations were run with an attacker fraction of 0.1.

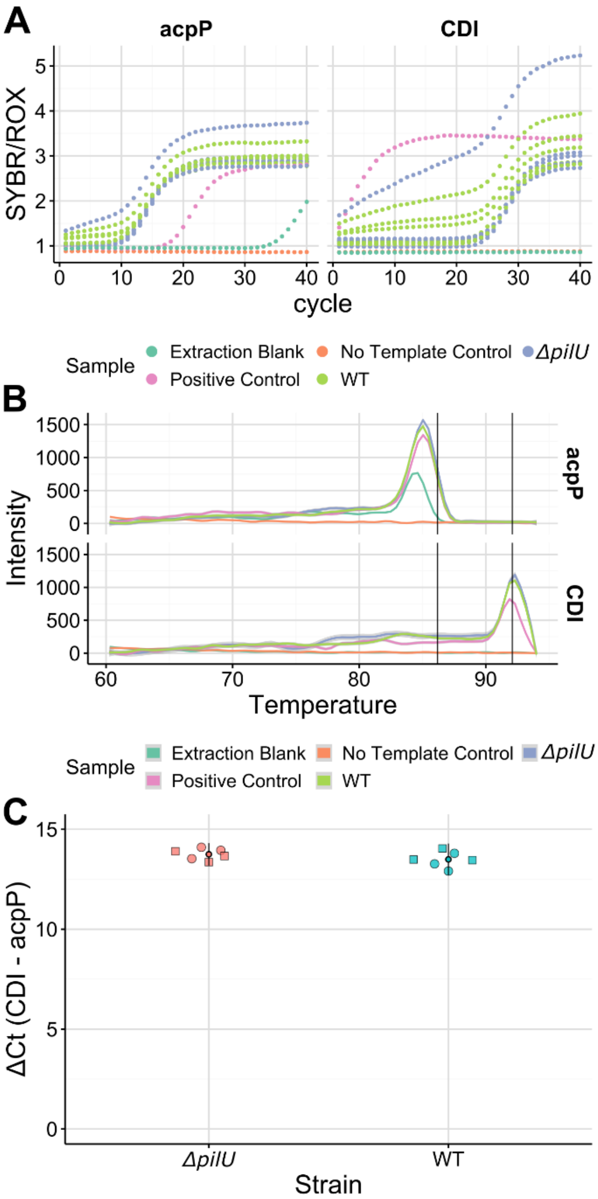

**Figure S5 | CDI expression is not altered in pilus mutants.** Quantitative reverse-transcriptase (qRT-PCR) of RNA extracted from colonies of wild-type and ΔpilU strains of *Pseudomonas aeruginosa* PAO1 shows that expression of CDI 1 (PA0040-PA0041) similar. **A** raw outputs (SYBR/ROX signal) for the house-keeping gene *acpP* and CDI, for samples including RNA extraction blank, qPCR control reaction with no template and gDNA (Positive Control). **B** Melting curves of products, with the predicted melting points (Tm) for *acpP* (86.2 C) and CDI (92.1 C). **C** ΔCt for wild-type and pilU deletion mutants. Each point is a single technical replicate qPCR reaction, squares and circles indicate distinct biological replicates. Data were analyzed in MXPro 4.1, see supplementary methods for details.

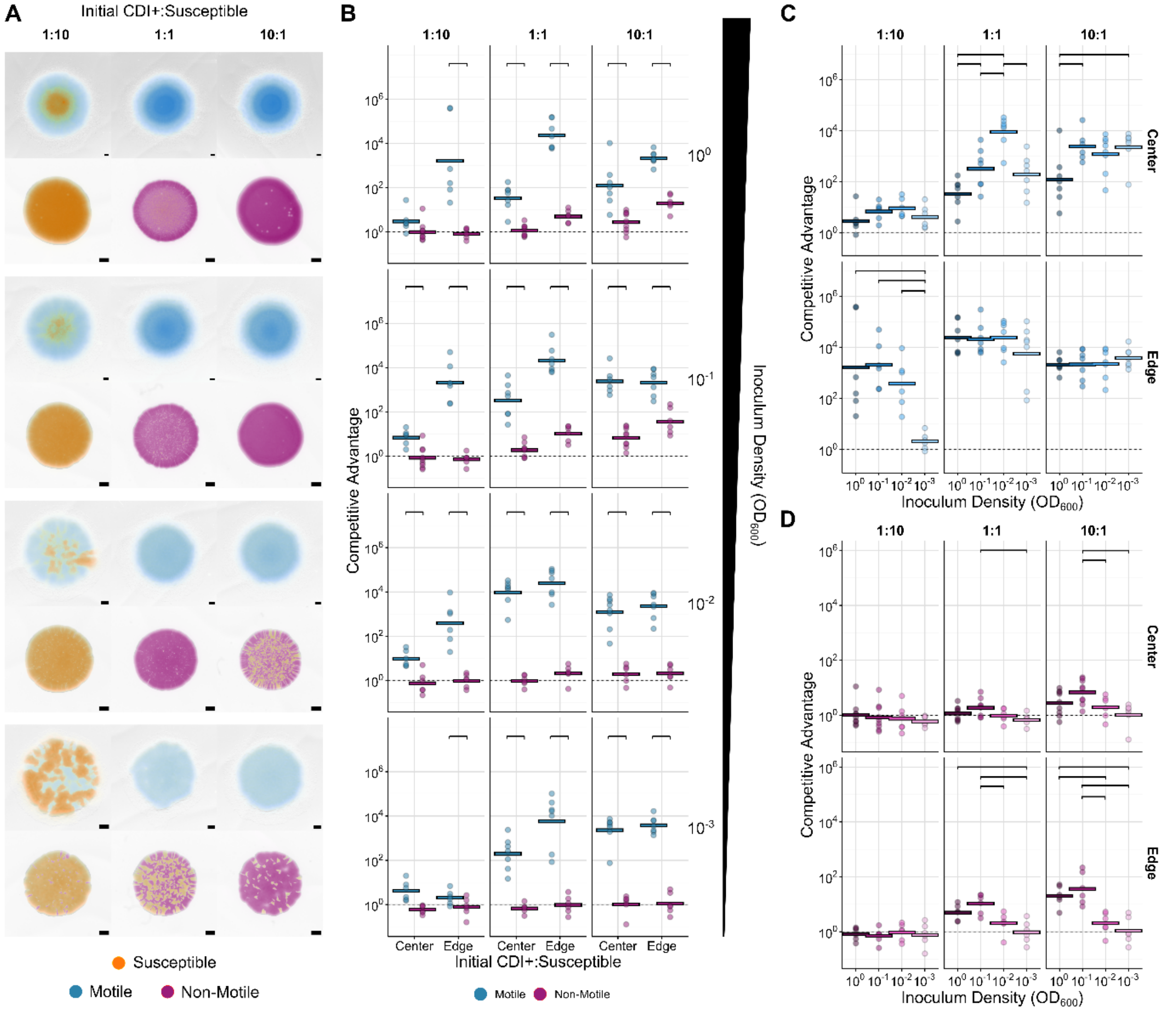

**Figure S6 | Motility enhances the competitive advantage provided by CDI at the center and edge of colonies regardless of the initial density and attacker frequency.** Colony competitions between wild-type or twitching motility deficient *P. aeruginosa* PAO1 and corresponding CDI-sensitve mutants were inoculated with different initial densities ($OD_{600}$) and ratios of attacker to sensitve cells. Representative microscopy images from competitions after 48 h of growth with motility active (top) and motility inactivated (bottom) (**A**) show differences in the scale and structure of communities. Quantification of the outcome of colony competitions picked at either the colony center or colony edge (**B**) reveal a consistently large advantage for motile attackers. Quantification of motile competitions from a range of inoculum densities and attacker:sensitve ratios (**C**) demonstrate a non-monotonic relationship between inoculum density and competitive advantage for 1:1 competitions at the colony center. Corresponding competitions using non-motile strains (**D**) do not show this effect. In **A**, strains are false-coloured either blue (motile CDI attacker), red (non-motile attacker) or yellow (motile and non-motile sensitive, top and bottom). Scale bar: 500 µm. Competitive advantage is calculated as the log fold-change in ratio of attacker:sensitve cells (as counted from sampling, plating and counting colony forming units) from the beginning to end of the experiment. Lines indicate the mean of replicates (n ≥

6). Top brackets indicate a significant difference between densities (one-sided Welch's t-test, $p < 0.05$, Benjamini-Hochberg MHT corrected 0.95).

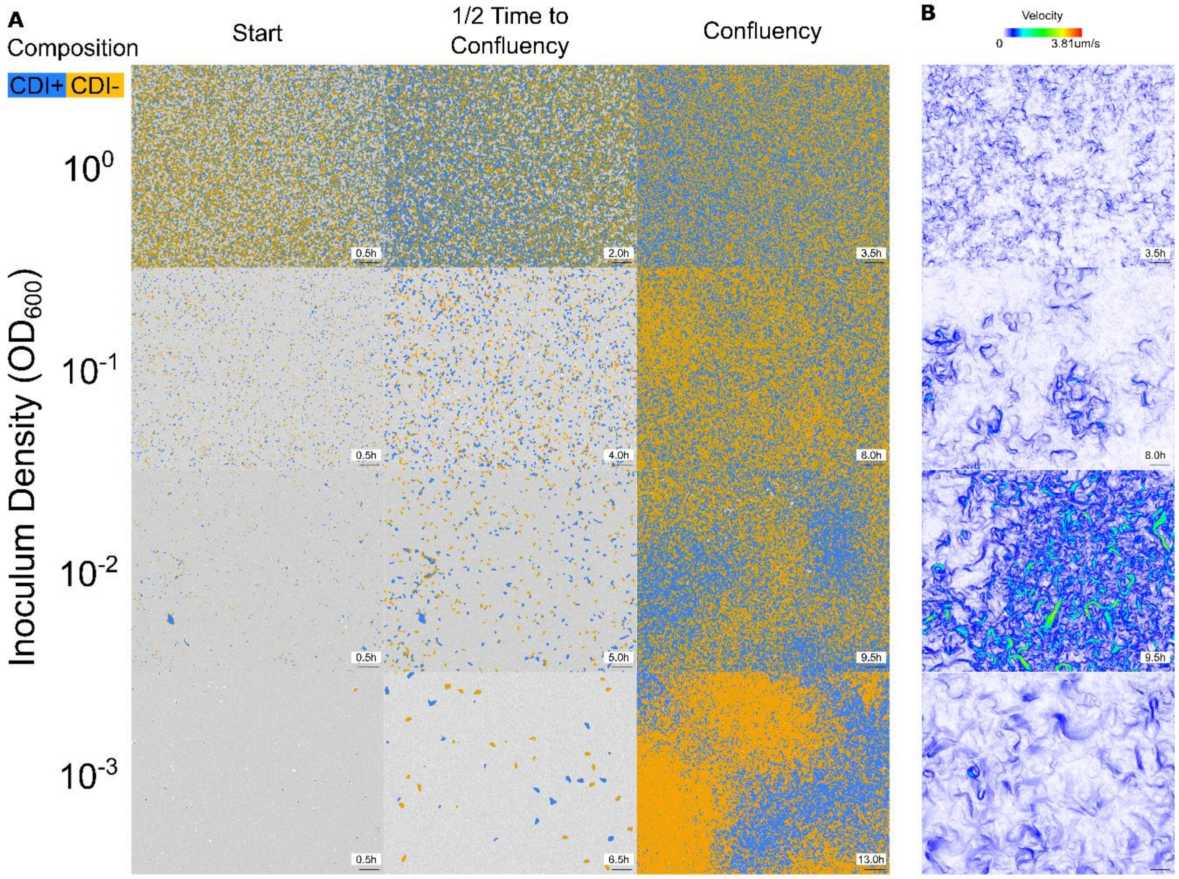

**Figure S7 | Full field of view images of timelapse and fluorescence microscopy showing differences of strain mixing and motility in colonies inoculated at different densities.** Colony competitions (1:1) between wild-type and CDI sensitive mutants were inoculated at different initial densities ($OD_{600}$) and imaged over time. Every 0.5 h after inoculation, a 1 min brightfield video was taken (0.5 frames $s^{-1}$) along with a single fluorescent snapshot in the YFP and mScarlet channels. Representative snapshots of colonies at the first time point ("Start", 0.5 h), the time when the surface was completely covered by cells ("Confluency", variable times), and halfway between the two timepoints ("1/2 Time to Confluency", variable times) (**A**) reveal increasing surface coverage with time, as well as the changing spatial distribution of wild-type attacker (blue) and CDI-sensitve (orange) strains. The velocity fields of colonies at confluency (**B**) also suggest an inverse relationship between inoculum density and cell motility. Images have been thresholded for display.

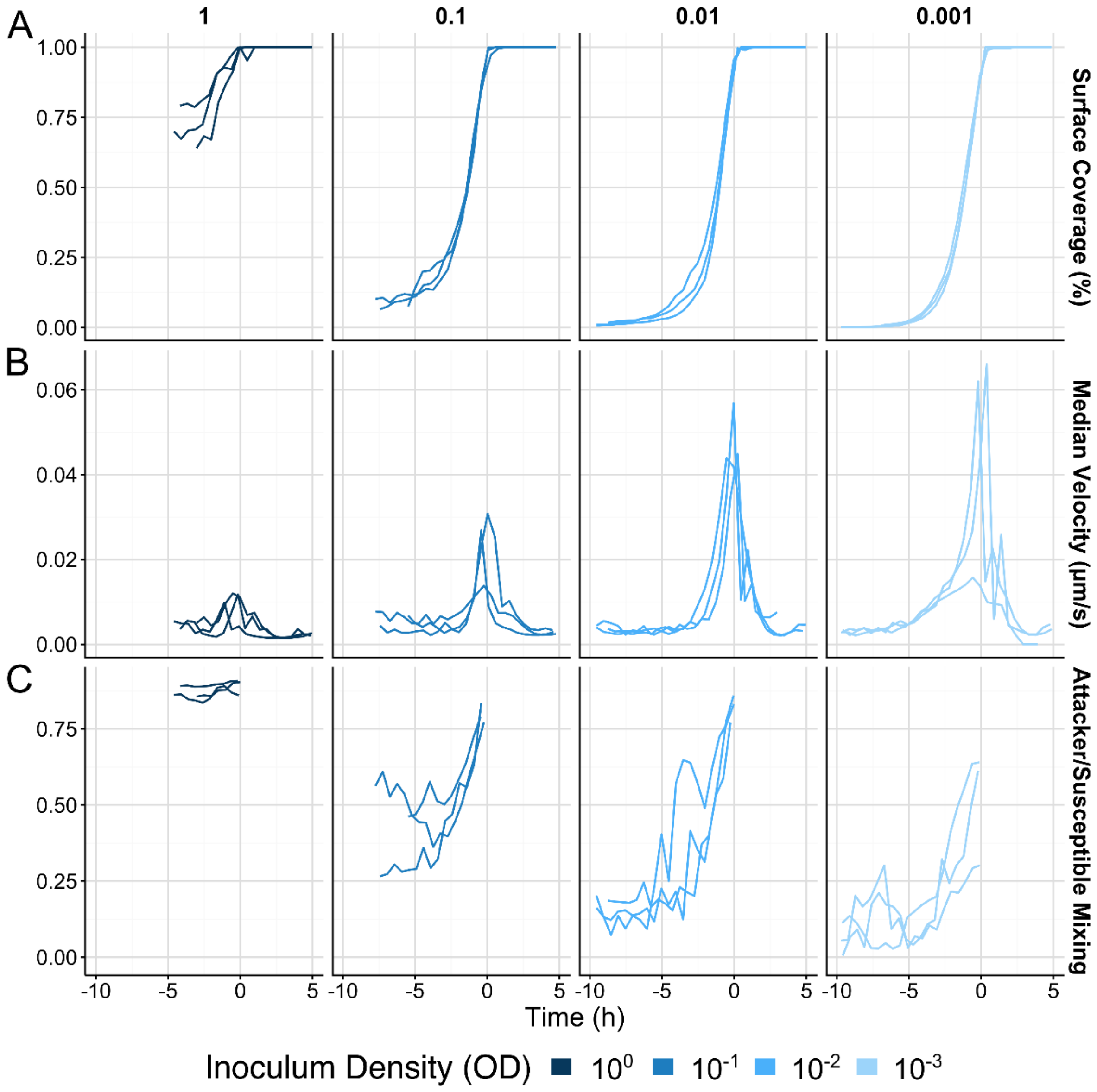

**Figure S8 | Individual replicate measurements of surface coverage, velocity and strain mixing from timelapse and fluorescent microscopy.** Individual replicates of colony competitions tracked in Figure 3. (**A**) percent area covered by cells, (**B**) median cell velocity magnitude and (**C**) extent of genotypic mixing between strains. Timecourses have been centered around the confluency time for each individual replicate.

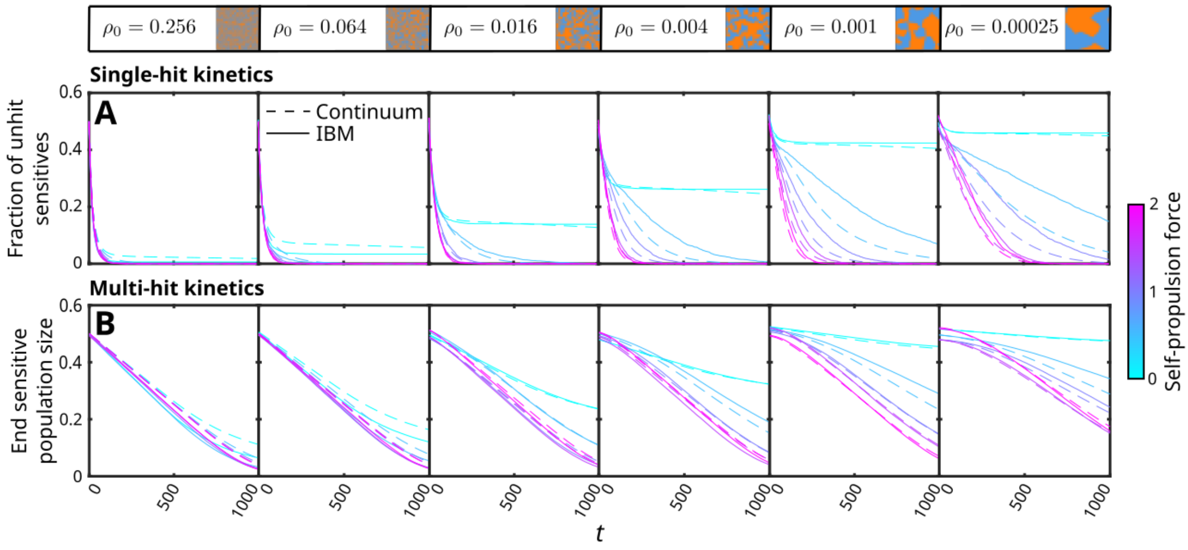

**Figure S9 | Matching between the IBM and the continuum model is robust to changes in inoculum ratio and toxin potency.** In our competition experiments, we consider colonies in which the attacker:sensitive inoculum ratio is 1:1 and the attacker's CDI toxin acts to slow down the growth of sensitives, rather than killing cells directly. To ensure that our matching between the two theoretical models was still effective with these changes, we performed additional simulations under these assumptions. In (**A**), we show the fraction of all cells that are sensitives that have not been hit as a function of time, equivalent to Fig. 1F but with a 1:1 rather than 1:9 inoculum ratio. This illustrates the killing dynamics under the assumption that one hit is sufficient to induce cell death. To simulate toxins that instead slow down growth with an accumulating impact with increasing numbers of hits, we introduce a toxin efficacy parameter $\xi = 0.02$ that represents the fractional slowdown in sensitive growth induced by each hit. In (**B**), we show the result of reducing the sensitive population size by the number of hits accumulated by each cell multiplied by $\xi$, which we take as a proxy for the final sensitive population size at the end of a competition experiment (Supplementary Note 4). Different seeding densities $\rho_0$ are indicated above each pair of plots, along with images of example continuum model starting states.

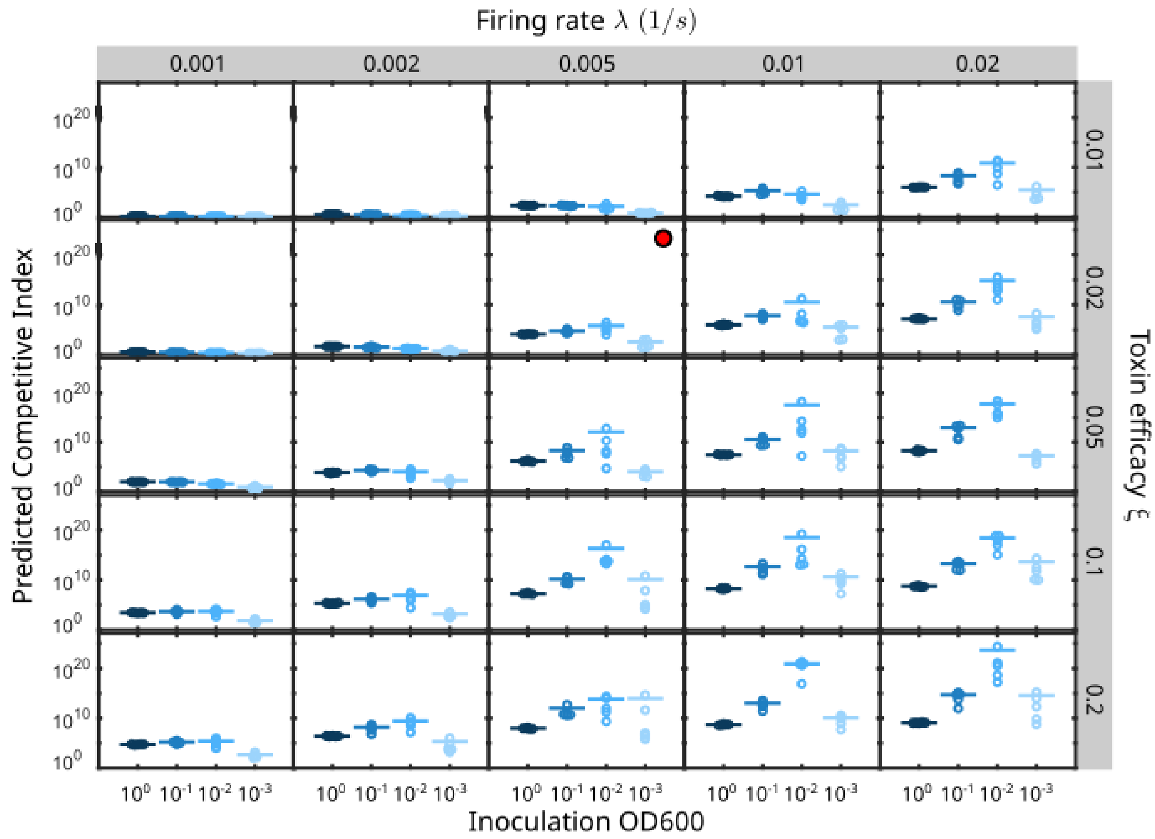

**Figure S10 | The predicted non-monotonic relationship between inoculation density and intoxification efficiency is robust to variations in CDI firing rate and toxin potency.** To test the impact of variations of our unconstrained parameters $\lambda$ and $\xi$ on our experimentally parameterized simulations (Fig. 4D-G), we ran a parameter sweep over these parameters while keeping all other factors fixed. Shown are the predicted competitive index of the attacker strain for *n*=5 simulations for each parameter combination, formatted equivalently to Fig. 4G. The combination of parameter values used in Fig. 4 is indicated with the red circle.

### Supplementary Movie Captions

#### Supplementary Movie 1 | IBM simulations of contact-dependent warfare demonstrate two mechanisms by which motility enhances intoxification.

We initialised IBM simulations with either initial genotypic patchiness (Patch +) or homogeneity (Patch -), and either with active motility of all cells (Motility +) or with no motility (Motility -). Attackers (blue) intoxify contacted sensitive cells (orange), with the number of hits accumulated by each sensitive cell indicated by the shade of orange (zero hits, dark orange, to 5+ hits, light orange). In each case, the simulated domain size is 200 x 200 cell widths, and the simulation is run for 1000 time units.

#### Supplementary Movie 2 | Continuum simulations capture large-scale intoxification dynamics.

The dynamics of the continuum model are illustrated with an example simulation. We simulate genotypic mixing as a diffusion process acting on an initial population distribution of attacker (blue) and sensitive (orange) cells, represented using a coarse-grained phase field (centre). Each lattice site contains a population of attackers and a distribution of different sensitive populations, varying by the number of accumulated contact-dependent hits as well as the current number of contacts to attackers. We represent this population structure for two example lattice sites in the phase field (purple and turquoise squares) using the stacked bar charts on the left and right. The population structure of sensitive cells is arranged as in Fig. 1E, with varying numbers of attacker contacts represented on the x-axis and increasing numbers of accumulated hits represented by the different colours of the bar sections (zero hits, dark orange, to 5+ hits, light orange). In this case, the initial seeding density $\rho_0 = 0.001$ cells $l_c{}^{-2}$, mean cell velocity $\bar{v} = 0.1\ l_c\ s^{-1}$, hit rate $\lambda = 0.01\ s^{-1}$, global fraction of attackers = 0.1 and the total simulation time was 1000 $s$.

#### Supplementary Movie 3 | Pilus-mediated motility in *P. aeruginosa* drives contact switching.

Observation of cells from an agar-surface colony shows how the direct contacts of a focal cell change over a short time period. A timelapse video was taken with brightfield images every 2 s, and fluorescent images every 32 s. On the left side, brightfield images show the positions of cells. Periodically the fluorescent image is shown and the video paused, allowing attackers (blue) and contact-dependent inhibition (CDI) sensitive cells (orange) to be distinguished. A single focal attacker cell is highlighted (red outline). On the right side, thresholded and segmented versions of the fluorescent images are shown, allowing the identity of the cells contacting the focal cell (cyan) to be discerned (inset). Contacts with clonemate attackers (blue), and uniquely identified sensitive cells (shades of magenta/red/orange/yellow) were manually determined at each fluorescently imaged timepoint. This video thus demonstrates that *P. aeruginosa* engages in contact switching.

#### Supplementary Movie 4 | Colonies inoculated at different starting densities display different patterns of motility.

Representative timelapse videos of agar-surface colonies of *P. aeruginosa* attacker and contact-dependent inhibition (CDI) sensitive cells inoculated at four different initial densities. Timelapses imaged every 2 s for 1 minute, every 30 min have been concatenated, of colonies inoculated from an initial density of OD600 1.0 (top left), 0.1 (top right) 0.01 (bottom left), and 0.001 (bottom right). Bacterial cells (white), grow, divide, and crawl across the agar surface (blue/black). The amount of motility increases as the density of cells on the surface increases, reaching a peak just before all available empty space on the surface is occupied by cells (confluency is reached), whereby then drops precipitously. Confluency is reached later for each lower inoculum density. Uncropped versions of these and similar timelapses were used to compute the velocity of cells over time. Time stamp is hours:minutes:seconds. Scale bar 10 µm.

#### Supplementary Movie 5 | Genotypic mixing varies as a function of inoculation density.

Representative fluorescent timelapse videos of agar-surface colonies of *P. aeruginosa* attacker and contact-dependent inhibition (CDI) sensitive cells inoculated at four different initial densities. Fluorescent snapshots were taken every 30 min of colonies inoculated from an initial density of OD600 1.0 (top left), 0.1 (top right) 0.01 (bottom left), and 0.001 (bottom right). Attacker (blue) and CDI-sensitive (orange) cells grow and divide until the agar surface (white) is completely covered (confluency). Once confluency is reached for each density, the image for this timepoint is held to allow comparison between densities. These and similar timelapses were used to quantify genotypic mixing. Time stamps are hours:minutes. Scale bar 100 µm.
